# A dual-photoreactive adapter enables one-step DNA barcoding of antibody panels for multiplexed imaging

**DOI:** 10.64898/2026.06.16.730979

**Authors:** Valeriy Pak, Yulia Ermakova, Christina Schniederjohann, Berkan Kanmaz, Robert Reinhardt, Felix Schneider, Tomas Martak, Sascha Dietrich, Peter-Martin Bruch, Sinem K. Saka

**Author notes:** These authors contributed equally (Valeriy Pak, Yulia Ermakova).

## Abstract

DNA-barcoded antibodies are central to a broad range of spatial and dissociative assays and applications, including multiplexed imaging, single-cell profiling, and proximity detection. Direct modification of primary antibodies with defined DNA barcodes enables flexible panel design for multiplexed labeling of proteins. However, conventional antibody-oligonucleotide conjugation methods are inefficient, low-throughput, and prone to batch variability, limiting the reliable generation of orthogonally barcoded antibody panels. These challenges are particularly acute during initial panel development, where lengthy protocols, conjugation failures, large antibody input requirements, and the need for custom antibody formulations increase experimental cost and effort. Site-specific conjugation strategies based on antibody Fc-domain binders offer a promising alternative. We streamlined this foundational approach to establish its compatibility with multiplexed imaging in cells and tissues; however, generating a full barcoded library is still resource intensive. This is because every unique DNA barcode must first be chemically linked to a separate binder before it can be attached to an antibody.

To overcome the prominent bottleneck of rapidly and reliably generating DNA-barcoded antibody panels, we introduce the universal oligo adapter (UnO) strategy. UnO fundamentally changes the workflow from barcode-specific conjugation to universal barcode conversion. We build on the established photoreactive protein G binder and combine it with a universal oligonucleotide carrying a second ultrafast photocrosslinking group, 3-cyanovinylcarbazole (cnvK). This creates a dual-functional adapter: one photoreactive group enables covalent attachment to the antibody, while the cnvK-containing universal oligo simultaneously captures a user-defined DNA barcode through hybridization and UV crosslinking in a single step. Rather than preparing separate conjugation reactions for dozens of barcodes, UnO acts as a single reagent that covalently couples any desired barcode onto small quantities of off-the-shelf primary antibodies in minutes.

We validate the generalizability and modularity of this approach across subcellular Immuno-SABER and tissue-based CODEX workflows for multiplex immunostaining. By converting antibody barcoding into a modular, one-step nucleic-acid adapter workflow, UnO reduces the cost, time, and complexity of generating DNA-barcoded antibody panels and provides an efficient, accessible solution to a central bottleneck in DNA-enabled multiplexed protein detection.

## INTRODUCTION

Affinity probe, particularly antibody, based identification of proteins via imaging remains the most widely utilized approach used for spatial protein detection and is almost unanimously employed for the new multiplexed protein imaging methods^1^, as well as in dissociative methods such as CITE-Seq and its derivatives^2,3^.

With the rise of the single-cell and spatial omics methodologies, the demand for multiplexing has been increasing, creating a need for direct labeling of primary antibodies (with fluorophores, DNA barcodes, metal polymers) to support large antibody panels. Oligo labeling of antibodies not only allows fast multiplexed imaging through cyclic hybridization of orthogonal fluorescence readout oligos^4–6^, but also enables additional capabilities such as in situ sequencing^7^ or next-generation sequencing readouts^2,3,8,9^, DNA-mediated signal amplification^6,10–15^, super-resolution imaging^16,17^, in situ barcoding^8,18^, or proximity detection^19–23^.

Yet, the direct labeling process for primary antibodies is hard to streamline, as each antibody is a unique protein and predominantly of unknown sequence. Conventional methods utilize bifunctional crosslinkers^24^, most frequently targeting antibodies on primary amines^25–27^ or sulfhydryl groups^28^. These conjugation approaches are however characterized with high antibody dropout rate and batch variation, low regiospecificity and throughput, and additionally require custom antibody formulations as well as multiple clean-up steps to remove unreacted/excess molecules making the conjugation process tedious, long, expensive and of low yield^1,29^. The glycosyl groups have also been successfully exploited for enzymatic site-specific labeling by transglutaminase^30–32^, though the enzymatic approach is relatively slow and the sugar groups may not be consistently present on all IgG isotypes.

As an alternative to direct modification of antibodies, a handful of methods have been developed in recent years, which take advantage of high-affinity binders such as protein G, protein A or anti-IgG nanobodies for functionalization of primary antibodies^17,33–37^. These binders typically recognize the Fc domain of antibodies and hence offer a generalizable site-specific labeling capability that is outside of the antibody paratope. Some of these binders have been further modified with photoreactive moieties, for example by introduction of the unnatural amino acid *p*-benzoylphenylalanine (BpA) outside of the Fc binding domain, to enable stable covalent linkage of the binder to the antibody^38–42^.

Here, we first build on these approaches and demonstrate the use of photocrosslinkable antibody binders for multiplexed immunostaining and signal amplification. Then we aim to increase the efficiency of creating barcoded antibody libraries by generating a universal oligo adapter (UnO). For this aim, we combine the antibody binder approach by functionalizing a photoreactive *para*-(4-hydroxybenzoyl)phenylalanine-substituted protein G fragment (here referred to as pG)^38^ with a universal oligonucleotide that is modified with an ultrafast photocrosslinking 3-cyanovinylcarbazole residue (cnvK)^43^. Thus, UnO consists of two photoreactive moieties, one for antibody-crosslinking (the pG) and one for DNA-interstrand crosslinking (the cnvK-bearing oligonucleotide), which effectively reduce the antibody conjugation procedure to simple irradiation of mixture containing primary antibody and DNA-barcode of interest with mild UV-light (365 nm, 6 min). Hence, UnO allows low-cost, high-efficiency, one-step antibody barcoding and enables going from small amounts of off-the-shelf antibodies to custom DNA barcoded libraries in a high-throughput manner within a couple of hours.

## RESULTS

Previously, recombinant photoreactive protein G, which bears non-natural amino acid benzoyl-phenylalanine for photocrosslinking, was used for labeling of antibodies in biochemical demonstrations^40^ and for live-cell labeling, followed by flow-cytometric analysis or super-resolution imaging^38^. We have built on these earlier studies to further develop this binder as a robust conjugation method for multiplexed antibody detection, particularly focusing on spatial proteomics (specifically multiplexed IF) applications, for which DNA-barcoded antibodies are in high demand both for custom (for example: DEI^44^, DNA-PAINT^17,45^/PRISM^46^, Immuno-SABER^6,14,47^, Seq-FISH+^48^, Cycle-HCR^49^, Spatial CITE-Seq^2^) and commercial assays (such as Akoya CODEX, Bruker/Nanostring CosMx^50^, 10X Xenium platforms, Ultivue/Vizgen InSituPlex^51^), and antibody functionality and specificity in a complex panel can be evaluated with relative ease by imaging of cells and tissues.

As the first step, we streamlined the protocol for production and purification of pG^38^ (**Fig. 1A**). We then optimized antibody photocrosslinking by increasing the pG to oligonucleotide ratio and improving the yield and quality by (i) removal of excess oligonucleotides prior to antibody conjugation, and (ii) removal of unreacted components after conjugation (see **Methods** for details of the protocols). In these experiments, we first conjugated the pG binder to 3 orthogonal 34 nucleotide long single-stranded DNA barcodes (**Supplementary Table 2)** carrying a terminal amino group and desalted the product. Then, we individually performed photocrosslinking of each pG-barcode to the antibodies targeting CD3, CD8 and CD68, followed by purification of the product using diafiltration columns. We confirmed high (nearly 100%) antibody conjugation efficiency by SDS-PAGE with Krypton and SYBR Gold staining (**Fig. 1B** and **Supplementary Fig. S1**).

**Figure 1:**
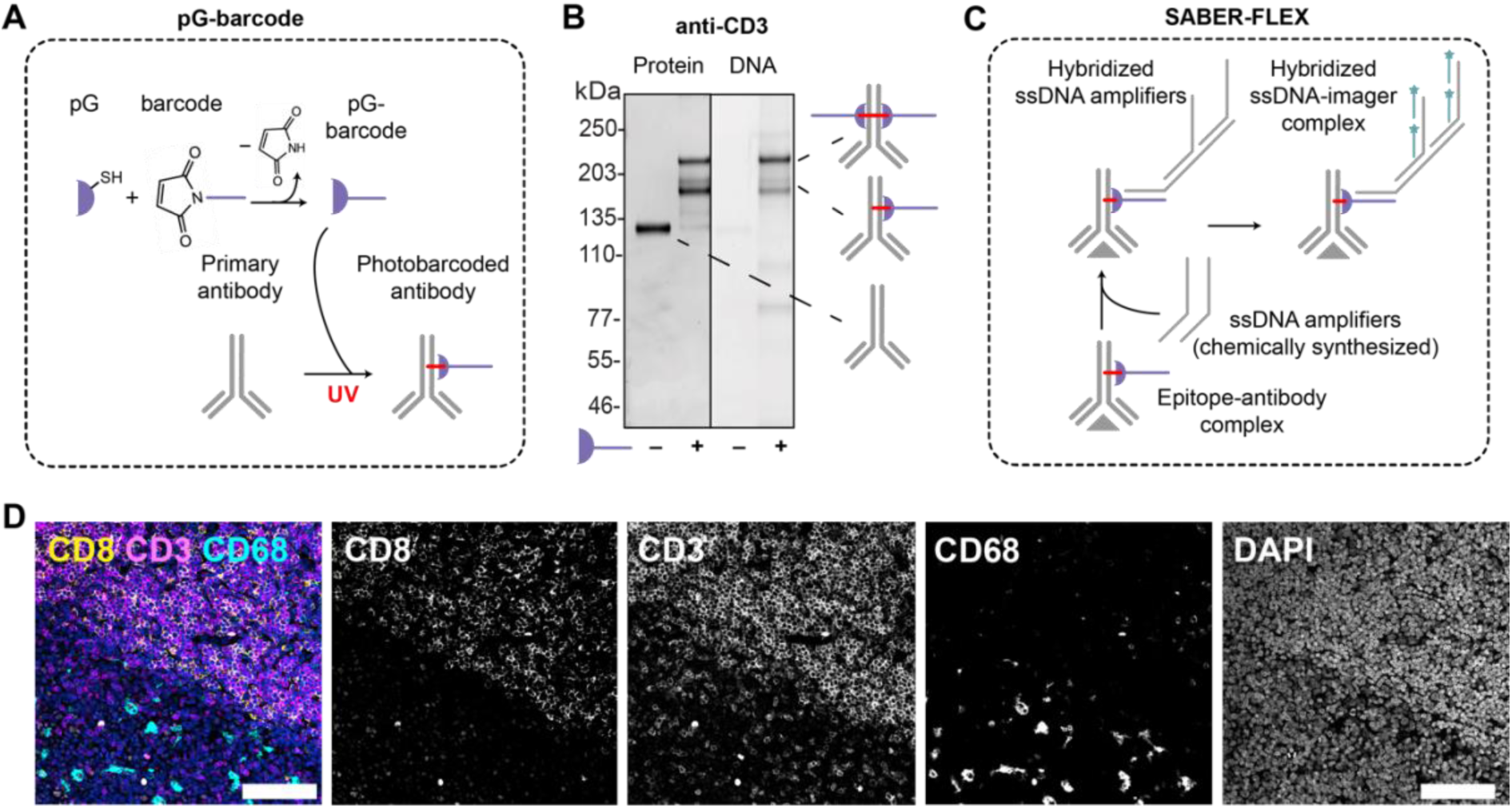
Combination of regiospecific binder-mediated antibody conjugation and orthogonal ssDNA signal amplifiers for multiplexed IF. (**A**) Scheme of antibody barcoding with protein G, shown in purple^38^. Protein G is conjugated to a DNA barcode via sulfo-SMCC crosslinker (only maleimide group is shown). The primary antibody is incubated with the resulting p G-barcode and illuminated with UV (365 nm, 6 min). (**B**) Representative non-reducing PAGE gel for barcoding of anti-CD3 rabbit monoclonal antibody. (**C**) SABER-Flex adapts ssDNA amplifiers from Xia et al. 2019^53^ for tunable signal amplification using chemically synthesized oligonucleotides. In contrast to PER concatemers, synthetic ssDNA amplifiers are typically limited by a maximum oligonucleotide synthesis length of ∼200 bases, therefore we used a branched detection scheme with secondary branched DNA amplifiers. (**D**) Maximum intensity projection of 3-plex SABER-Flex staining of human FFPE tonsil section (5 µm thick) with DNA barcoded rabbit antibodies targeting CD8, CD3 and CD68. Scale bar is 100 µm.

For assessing the functionality of conjugates, we utilized multiplexed IF, where we performed simultaneous branched DNA signal amplification by use of single-stranded DNA (ssDNA) concatemers and detection by fluorescent oligonucleotides (“imagers”). The concatemeric

DNA amplifiers can be generated either by primer exchange reaction^52^ as we previously showed in our Immuno-SABER approach^6^) or alternatively by a more flexible adaptation of the method, which we refer to as SABER-Flex (**Fig. 1C**). In this case, PER-generated concatemers are replaced with synthetic oligonucleotides that could be commercially sourced (as has also been shown before for RNA ^53^) (see **Supplementary Table 1** for comparison between Immuno-SABER and SABER-Flex). We used SABER-Flex for 3-plex imaging with protein G-barcoded antibodies (CD3, CD8, CD68) in 5 µm FFPE human tonsil tissue (**Fig. 1D**). All conjugated antibodies showed high conjugation efficiency **(Supplementary Fig. S1)** and highly specific labeling of the target cell types in FFPE human tonsil tissues (all T-cells, cytotoxic T-cells and macrophages, respectively for CD3, CD8 and CD68), validating intact functionality and absence of background and visible crosstalk after the conjugation and mixing of conjugates for simultaneous staining (**Fig. 1D**, **Supplementary Fig. S2-S3**).

This initial characterization demonstrates reliable application of the pG barcoding for multiplexed IF. However, in these cases, a separate conjugation to pG must be performed for each DNA barcode first, and then these barcoded products are individually used for site - specific antibody conjugation with photocrosslinking. Thus, DNA-conjugation of the antibody binder limits the throughput, especially as the plexity increases. The need to generate a separate conjugated product for each barcode also renders purification challenging and low yield resulting in reduced final quality. To address this problem, we set out to generalize the binder-mediated barcoding strategy and designed a bifunctional adapter, by incorporating a universal oligonucleotide that also carries a photocrosslinkable domain, enabling one-step modification of conjugates with desired barcode oligos.

### Universal oligo adapter (UnO) design

For the oligo photocrosslinking moiety, we chose 3-cyanovinylcarbazole (cnvK), which is an ultrafast photoreactive base analog that enables UV-triggered, site-specific crosslinking of oligonucleotides to complementary strands^18,43,54^. Hence, UnO carries two photoreactive moieties, one for antibody-crosslinking and one for DNA-interstrand crosslinking, which can be both activated simultaneously for one-step conjugation via mild UV light at 365 nm. We have designed a universal DNA sequence to hybridize to a constant complementary part of the barcodes to ensure efficient hybridization at room temperature and a balanced GC-content and lack of secondary structure and homopolymer regions (see **Methods** for details), as well as low similarity to frequently used multiplexing barcode sequences in the literature (Immuno-SABER^6^, CODEX^28^ and CITE-seq^2^). We have generated an efficient FPLC-free protocol with DNA-coated magnetic beads for high-purity bench-top production of UnO (**Fig. 2A-C**). cnvK modified oligos can be produced by in-house chemical synthesis or can be sourced from DNA synthesis companies. To keep the costs for the modified oligo minimal, we aimed to achieve high efficiency by saturating the reaction with an excess of the recombinant binder **(Fig. 2A**).

**Figure 2:**
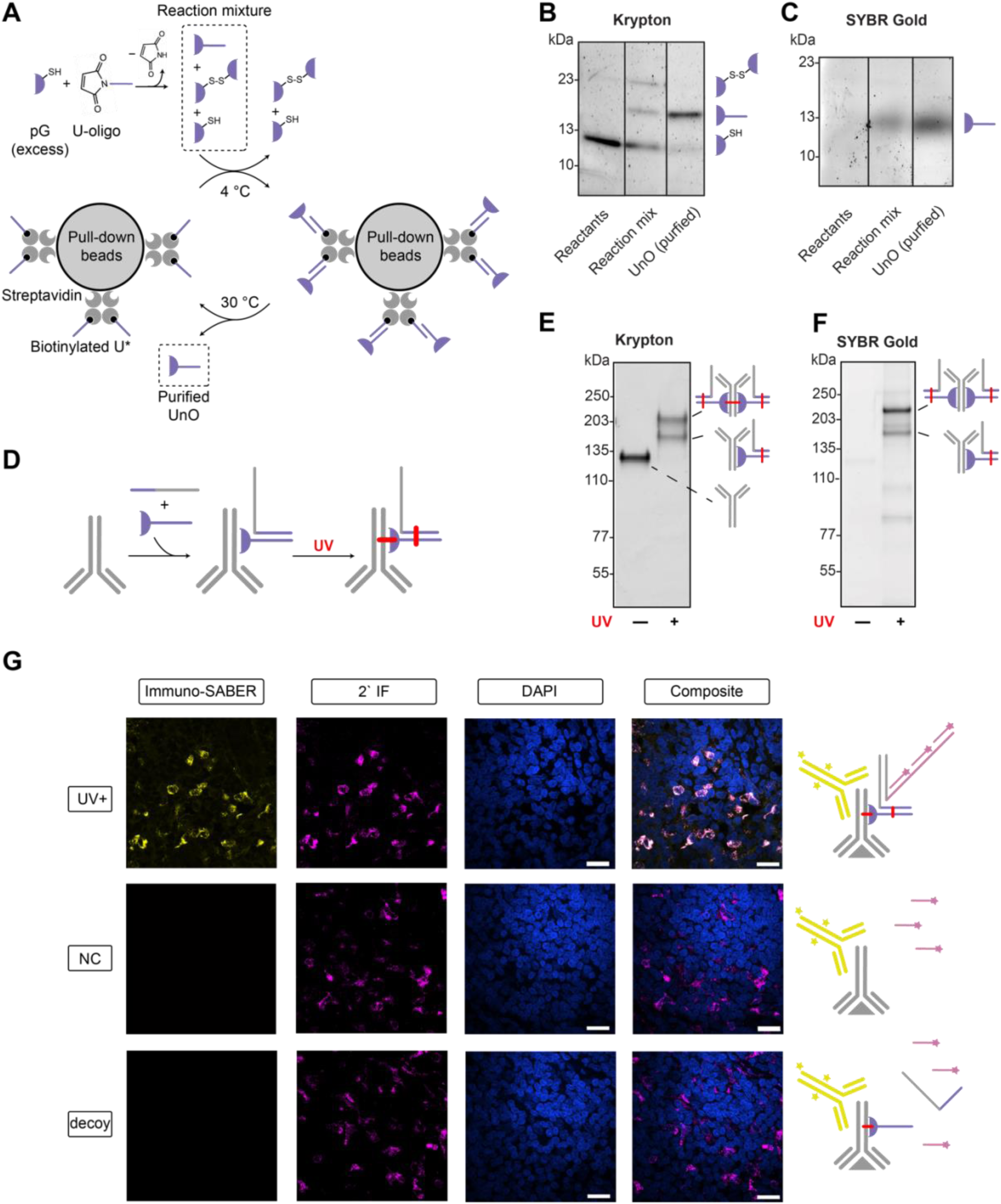
UnO production and antibody barcoding. (**A**) UnO is synthesized by coupling of pG with an excess of cnvK-containing universal adapter oligo (U-oligo). The resulting mixture of UnO and unreacted excess of pG is purified via pull-down using streptavidin beads functionalized with a complementary 11-mer capture oligonucleotide (U*-oligo). (**B**-**C**) SDS-PAGE gels demonstrating UnO synthesis and purification: reactants (left lanes), UnO reaction mix (middle lanes) and after purification (right lanes) for Protein (Krypton, **B**) and DNA (SYBR Gold, **C**) staining. Higher molecular weight product (∼23 kDa) represents the product of pG cysteine oxidation throughout the procedure. (**D**) Antibody barcoding scheme: antibody and oligonucleotide barcode (containing UnO complementary sequence, shown in violet) are mixed with UnO and illuminated with UV, leading to covalently linked (red) product in a single step. (**E**-**F**) Representative SDS-PAGE gels demonstrating primary antibody (anti-CD68) conjugation, with protein (Krypton, **E**) and DNA (SYBR Gold, **F**) staining. Two shifted bands represent single- and dual-labeled antibodies. Based on gel quantification (**Methods**), we observed the labeling efficiency to be ∼99% (n=2 technical replicates). (**G**) Fluorescent micrographs of 5 µm-thick human FFPE tonsil section stained with anti-CD68 antibody conjugated with UnO and irradiated with UV ("UV+", top row). Non-conjugated primary antibody (“NC”, middle row) was used as a positive control (conventional IF with secondary antibodies done on consecutive sections). As a negative control, pG conjugated to non-cnvK modified oligonucleotide with the same sequence as U was used to barcode the antibody ("decoy", bottom row). Primary antibody staining for SABER-Flex was performed in parallel to conventional indirect IF. Fluorescent channels (from left to right) correspond to: Immuno-SABER (ATTO-565), indirect IF (Alexa Fluor 635 conjugated secondary antibody), DAPI and composite image (pink and yellow pseudo-color representing Immuno-SABER and IF channels, respectively. White indicates colocalization between the two channels). Scale bar is 50 µm.

We then created a workflow for one-step antibody barcoding with UnO, where both photocrosslinking reactions happen in parallel to yield the site-specifically labeled antibodies with the desired barcodes (**Fig. 2D**). By applying long-wavelength UV exposure of ≤6 min (see **Methods**), we achieved labeling of ∼99% of antibody with the desired barcode oligo (**Fig. 2E-F**). We tested UnO-barcoded antibodies with Immuno-SABER staining of human tonsil sections, followed by conventional immunofluorescence with secondary antibodies (**Fig. 2G**). We observed identical staining patterns between the Immuno-SABER and indirect IF with barcoded or non-conjugated primaries (**Fig 2G**, bottom row - "NC"), demonstrating the functionality of the UnO barcoded antibodies. As an additional negative control, we barcoded the same antibodies with “decoy” UnO, which does not carry the cnvK modification, under the same illumination with UV, which produced no staining in Immuno-SABER channel, indicating cnvK-dependent mechanism of the photo-barcoding.

We then used the same scheme for a more quantitative evaluation of the performance of UnO-barcoded antibodies for higher resolution subcellular labeling experiments **(Fig. 3A, Supplementary Fig. S4,5)**. We labeled antibodies against α-tubulin, vimentin and GM130 (each having distinct subcellular localization pattern) with orthogonal Immuno-SABER barcodes (**Supplementary Tables 3,5**). We used them for single-target staining of fixed HeLa cells according to the Immuno-SABER protocol and detected the antibodies with (1) cognate imager oligonucleotides, and (2) secondary antibodies conjugated to a spectrally distinct fluorophore for colocalization analysis. As controls we used (1) non-conjugated and (2) non-crosslinked (i.e. the reaction includes the same steps except UV illumination) ("UV-" control) antibodies. We observed (1) similar staining patterns between intact and barcoded antibodies, (2) high Pearson correlation between Immuno-SABER and IF signals (for GM130: 0.93; alpha-tubulin: 0.92; vimentin: 0.97), and (3) no staining above background level for UV– control. Thus, UnO-barcoded antibodies are suitable for multiplexed protein detection.

**Figure 3:**
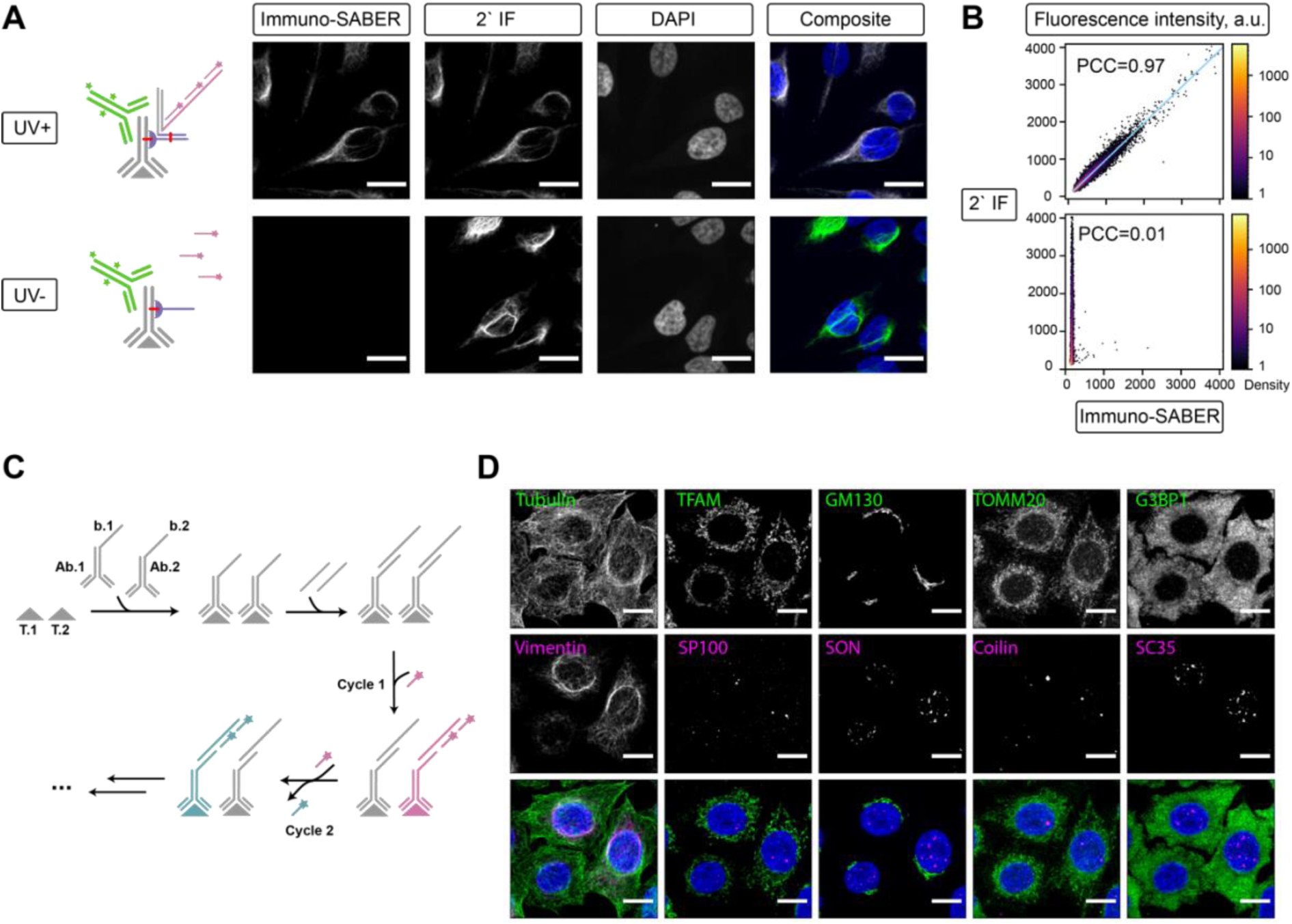
Application of UnO-barcoded antibody panels for multiplexed imaging. **(A)** Fluorescent micrographs of fixed HeLa cells stained with anti-vimentin antibody conjugated with UnO and irradiated with UV (“UV+”, top row) or non-irradiated control (“UV-“, bottom row). Primary antibody staining for Immuno-SABER was performed in parallel to conventional indirect IF. Fluorescent channels correspond to (from left to right): Immuno-SABER (ATTO-565), indirect IF (Alexa Fluor 635 conjugated secondary antibody), DAPI and composite image (pink and green pseudo-color representing Immuno-SABER and IF channels, respectively, with white showing colocalization between two channels). Scale bar is 20 μm. (**B**) Fluorescence intensity scatterplot (raw intensity units) for Immuno-SABER (x-axis) and indirect IF (y-axis) channels for the corresponding “UV+” and “UV-“ images in (**A**). Pearson correlation coefficients are indicated on the plots. Blue line represents linear regression fit. Individual points are colored by local normalized density (log-scale color bar is indicated on the right). (**C**) Schematic of multiplexed Immuno-SABER in cultured HeLa Kyoto cells schematics. (**D**) Fluorescent micrographs of fixed HeLa cells stained with the following UnO-barcoded antibodies: α-tubulin (microtubules), vimentin (intermediate filaments), TFAM and TOMM20 (mitochondria), GM130 (golgi), G3BP1 (stress granule marker; under non-stressed conditions distributed throughput the cytoplasm) and nuclear bodies (SP100, SON, coilin, SC35). DAPI counterstaining is shown on composite images in blue. Scale bar: 20 µm.

### Streamlined creation of custom panels for multiplexed IF

A common limitation of antibody DNA-labeling approaches is unpredictable labeling efficiency of antibodies with different isotypes or raised from different hosts due to amino acid sequence/tertiary structure differences. It is important to note that protein G, which is derived from *Streptococcal* bacteria, can natively bind to a broad range of IgGs at the CH2-CH3 junction^55^, with varying binding affinities depending on the IgG subtypes and host organisms. These affinities have been characterized by previous work, since such binder molecules are commonly utilized for affinity purification protocols and are part of many commercial products^56,57^. While pG has very high affinity towards rabbit IgG, human IgG1-4, and mouse IgG2a/b, the BPA moiety has been shown before to not crosslink well to antibody subtypes that lack methionine residues in the targeted binding site of the Fc-domain, including mouse IgG1 (mIgG1) antibodies^40^. Since this is a popular antibody subtype for immunostaining applications, we utilized an alternative binder for mIgG1 to show how different antibody binders can be turned into bifunctional reagents for the UnO approach. To this end, a commercially available mIgG1-targeting antibody binder with an azide modification (AlphaThera) was conjugated to the aminated *U-oligo* using TFP-DBCO as a bifunctional crosslinker (see **Methods**) and was utilized as an alternative UnO adapter termed mUnO (“mouse IgG1 UnO”).

While we have successfully demonstrated DNA-barcoding of mIgG1 isotype antibodies using mUnO (**Supplementary Fig. S6**), the overall antibody labeling efficiency for this adapter on the gels was lower than that of UnO. We hypothesize that this is due to lower labelling efficiency of the mIgG1 binder compared to pG, as antibody labelling failed to reach full saturation despite providing an excess of mUnO in the reaction mixture. Nevertheless, we were able to obtain a functional conjugate that is sufficiently performant for the downstream imaging applications. To validate the functionality of UnO-conjugated antibodies for multiplexed detection, we created a panel of 10 DNA-barcoded antibodies targeting major organelle markers for subcellular analysis (**Fig. 3D**). We conjugated the 10 primary antibodies of the panel (including 2 of mouse IgG1 isotype, conjugated with mUnO) to unique DNA barcodes (**Supplementary Fig. S6** and **Supplementary Tables 3,5**) and applied the panel to stain cultured cells with the Immuno-SABER protocol (see **Supplementary Tables 4** for PER reaction conditions). We profiled all 10 target epitopes within four cycles, with each cycle consisting of the following steps: (1) staining with a subset of 2-3 targets with cognate imager oligonucleotides, (2) imaging of multiple fields-of-view, and (3) dehybridization of imagers prior to the next staining round. (**Fig. 3C**).

All four cycles were performed sequentially on the same fields of view, which were computationally registered to align the same cells across cycles. The panel spans compartments with distinct staining morphologies, including filaments (α-tubulin, vimentin), puncta (SP100, coilin, SON, SC35) and cisternae (GM130), and all markers showed the expected subcellular distribution with high signal-to-background ratio **(Supplementary Fig. S7**).

To evaluate the panel as a whole rather than marker by marker, we clustered image pixels in marker-intensity space using self-organizing maps (see **Methods**). Pixel clustering in marker-intensity space groups pixels into recurrent marker combinations referred to as multiplexed cell units (MCUs), an approach introduced for subcellular protein maps in cultured cells^58^. The analysis is unsupervised and receives no information about marker identity, so the panel serves as its own benchmark: the antibodies were selected for well-characterized localizations, and recovering those compartments from the barcoded readout alone tests whether the conjugates report the expected spatial organization. Clustering resolved pixel groups corresponding to the major compartments sampled by the panel, including mitochondria (MCU7; TFAM, TOMM20) and Golgi (MCU9; GM130), and separated the nuclear speckle markers SON and SC35 (MCU8) from SP100 (MCU3) and coilin (MCU6), which mark typically adjacent but distinct nuclear bodies (**Supplementary Fig. S8**). Together, these experiments demonstrate that UnO-barcoded panels support rapid development of performant custom antibody panels for multiplexed immunofluorescence, with the resulting images recapitulating known subcellular organization.

To further demonstrate the broad applicability of UnO-barcoded antibodies in alternative DNA-based MxIF approaches, we designed an experiment combining the commercial CODEX protocol with a custom UnO-barcoded antibody panel. The CODEX protocol typically utilizes a conjugation approach which depends on partially reducing inter-chain disulfide bonds to create labile sulfhydryl groups used for coupling of maleimide-modified barcode oligos to cysteine residues^5,28,59^. We developed an antibody panel with off-the-shelf targeting common cell type markers found in human tonsil using UnO adapter with the same barcode sequences as in the standard CODEX assay (**Supplementary Table 6**) and used this panel to stain FFPE tonsil sections (**Supplementary Figures S9-S10**). In parallel, we conjugated the same antibody clones targeting the Cysteine groups according to the original CODEX protocol and stained an adjacent tissue section. Since this conjugation protocol cannot tolerate additives like BSA, antibodies were obtained from the vendors with custom formulations to allow cysteine-directed labeling. Both sections were then imaged on the PhenoCycler-Fusion 2.0 imaging system (Akoya Biosciences/Quanterix). Upon visual inspection and by quantification of marker positivity for each cell, we confirmed that UnO consistently performed *on par* with the cysteine-directed CODEX barcoding protocol, with markers exhibiting well-resolved, compartmentalized, and consistent staining patterns and cell type identifications across the tissue section, including effector and cytotoxic T lymphocytes (**Supplementary Fig. S9A-C**). Notably, CD38, a key marker for activated lymphocytes and plasma cells, was only detected in the tissue stained with UnO-barcoded antibodies, showing a strong and specific signal which was absent in the cysteine-directed conjugation protocol (**Supplementary Fig. S9C**). Antibody dropout after conjugation to cysteine residues is a recognized practical issue, which might be attributed to the relatively harsh reaction conditions used for partial reduction of antibody inter-chain disulfide bonds, leading to the destabilization of native tertiary structure and loss of functionality^29^. Thus, UnO is compatible with commercial multiplexing workflows like CODEX and has potential to decrease failure rates for DNA-barcoded antibodies.

Overall, we demonstrate that both pG and UnO support multiplexing with site-specific high efficiency DNA-barcoding. UnO can be used as a single reagent for one-step parallelized generation of panels of antibodies with different barcode sequences. Since UnO conjugation can be performed at scales down to <5 µg amounts, it is particularly suitable for initial panel establishment and characterization. Both the antibody and the oligo moiety of the conjugated products work reliably after the dual crosslinking reaction as the targets can be detected both with secondary antibodies and single-stranded oligos that hybridize to the barcodes (either imagers like in CODEX or amplifier sequences like in Immuno-SABER).

## DISCUSSION

UnO is a versatile universal adapter strategy designed to address a principal bottleneck in DNA-based multiplexed protein imaging: the efficient generation of large, quantitative, and reproducible antibody panels. Conventional panel assembly is frequently labor-intensive, time-consuming, expensive and prone to suboptimal yields, particularly when scaled to numerous targets. UnO mitigates these limitations by enabling small-scale antibody–oligonucleotide conjugations for off-the-shelf antibodies with a single universal reagent, thereby facilitating rapid pilot panel construction prior to large-scale implementation. This workflow minimizes antibody consumption, expedites the panel optimization process and supports better defined degree of labeling (up to 2 barcodes per antibody).

We have validated the suitability of UnO for DNA-based detection of antibodies across custom or commercial multiplexed imaging protocols and sample types (cultured cells, FFPE tissue sections), demonstrating compatibility with a diverse fixation, staining, and imaging workflows. In contrast to nanobody- or Fab-fragment–based approaches (e.g. SUM-PAINT^17^), which employ a non-covalent barcoding strategy, UnO minimizes the risk of the barcode crosstalk due to the formation of a stable covalent linkage between antibody and oligonucleotide. In contrast to nanobodies, the UnO system applies a single universal conjugation chemistry to any compatible antibody, enabling efficient expansion to large, multiplexed panels while preserving long-term stability of the label. The covalent bond formed between the universal oligonucleotide and the antibody confers stability, promoting barcode retention during stringent washing, prolonged storage, or long multi-cycle imaging protocols, preventing loss of signal and crosstalk between targets^60^.

UnO approach also allows a high flexibility with the DNA barcode sequence. Previously, it was shown that conjugation efficiency is substantially decreased when longer DNA barcodes are utilized for conjugation of oligos to Protein G binder^38^ or directly to antibodies^27^. UnO alleviates this limitation by using a short universal oligonucleotide during the conjugation step and achieving barcode conversion through hybridization followed by covalent crosslinking. This design maintains high conjugation efficiency while preserving flexibility in barcode sequence selection. In our experiments, we successfully used panels with a range of barcode oligonucleotide lengths (of up to 63 nucleotides) across different multiplexing workflows. It is worth noting that conjugation of both protein G and oligonucleotides confers additional negative charges, which may introduce potential ionic effects. We did not observe these to have a major impact on antibody conjugation or immunostaining performance in our experiments. However, due to the additional charge, the conjugated product is typically stained less efficiently on the PAGE gels than the unconjugated antibody, which may lead to an underestimation of conjugation efficiency, as discussed in the **Methods**.

With the exception of a few antibodies that we used initially for the evaluation of UnO efficiency (CD3, CD8, CD68), all our UnO conjugations utilized off-the-shelf antibodies (see **Supplementary Tables 5-6**) that are typically stored at lower concentration and in buffers containing stabilizers including sodium azide, BSA, glycerol, which would prevent application of most other conjugation approaches (for instance, partial reduction of disulfide bonds used for conjugation of antibodies for the CODEX workflow^5,28^). Among these additives BSA is typically the hardest one to remove from antibody stocks due to its high concentration and molecular weight. The pG fragment we use does not contain the albumin interacting domain and was shown not to cross-react with BSA^38^. It was also previously shown that BSA and Tween-20 were largely tolerated, but azide and glycerol may decrease the conjugation efficiency of *p*-Bpa^38,61^. Therefore, although we satisfactorily applied UnO conjugation for off-the-shelf antibodies, even higher efficiencies might be obtained with custom formulated antibodies, which are typically supplied at higher concentrations and without additives. We also noticed that antibodies purified using protein A or G by vendors, may carry leftover protein A/G in the stocks, which would potentially decrease the conjugation efficiency with pG or UnO.

For most applications, a simple ultrafiltration clean-up would be sufficient to remove excess barcodes and unconjugated UnO. For highly quantitative applications that are very sensitive to background or for specific antibodies and antibody types where conjugation efficiency is low, it might be desirable to purify the barcoded antibodies more specifically. For such cases, UnO offers the possibility of coupling DNA-based purification and barcode conjugation to achieve a more complete and precisely defined degree of labeling. We have previously utilized this kind of optional one step purification through DNA-based capture on beads and controlled release^6^ and it can be easily performed for tens of antibodies at the same time and at small scales.

The current implementation of UnO exhibits certain constraints. Use of pG for Fc-targeting restricts compatibility to a subset of antibody isotypes, potentially excluding some IgG subclasses or antibodies from less common host species. This is an inevitable limitation of using antibodies raised in various species which might be mitigated by further development of alternative antibody binders targeting specific isotypes. To exemplify this, we demonstrated modularity of UnO design by an alternative employing mouse IgG1-specific antibody binder to successfully barcode corresponding antibody clones and use them in mixed antibody panels. This experiment demonstrates the potential of UnO-approach for the development of isotype-specific bifunctional adapters with variable antibody-binding moiety (e.g., protein G^38^, protein A^62,63^, protein Z^64^, nanobodies^17^, FcIII peptide^65^ and others). We envision further development and wide application of binders, such as photoreactive versions of secondary nanobodies, would be greatly useful for creating custom panels of virtually unlimited multiplexity in a high-throughput fashion.

In summary, UnO offers a customizable universal adapter approach to address a major bottleneck in generating DNA-barcoded antibody panels and offers improvements in cost, fidelity, throughput and ease of panel creation for multiplexed protein detection. Beyond multiplexed immunofluorescence imaging, the universal adapter concept underlying UnO has potential applications in CITE-seq, for modular attachment of sequencing-compatible barcodes, in proximity-based detection assays such as the proximity ligation or proximity extension assays, and in spatial multi-omics platforms, where the coordinated mapping of proteins and nucleic acids within the same tissue section is desirable.

## MATERIAL AND METHODS

### Recombinant pG expression and purification

E. coli BL21 (DE3) were transformed with equimolar mix of pET28a-pG (encoding protein G with amber stop-codon at the unnatural amino acid position 38, modified with N-end Strep-tag and C-tag His-tag) and pEVOL (encoding unnatural amino acid tRNA and tRNA-synthase) plasmids (both kindly provided by Glenn Cremers, de Greef Lab, Eindhoven University). Bacterial culture was incubated in 20 ml of LB medium supplemented with 50 µg/mL kanamycin and 25 µg/mL chloramphenicol overnight at 37 °C. Overnight culture was used to inoculate 2 L of 2xYT medium supplemented with kanamycin and chloramphenicol (in 5 L flasks, 10 mL of preculture per 1 L). Cultures were grown until OD600 = 0.8 and then supplemented with 1 mM para-(4-hydroxybenzoyl)-phenylalanine and induced with 1 mM IPTG and 0.02% arabinose. Cultures were incubated overnight at 18 °C. Cells were harvested and the pellet was stored at −20 °C before isolation.

Cell pellet (from 2 L of culture) was resuspended in 50 mL of lysis buffer (1xPBS, 20 mM imidazole, 370 mM NaCl, 10% glycerol). The cells were lysed by sonication (8 × 45 seconds with 1 minute break in between rounds on ice). Supernatant was centrifuged at 35,000 rpm for 30 min at 4 °C (Ti45 rotor, Beckmann UC) and filtered. The filtered supernatant was loaded on a 50 mL HisTrap column (GE, equilibrated in lysis buffer) with 0.5 mL/min flow. The column was washed with the lysis buffer until a stable baseline was reached. The protein was eluted with a step gradient to 100% elution buffer (1xPBS, 250 mM imidazole, 370 mM NaCl, 10% glycerol). The elution fraction was directly loaded onto the 5 mL Strep-Tactin column equilibrated in strep wash buffer (100 mM Tris-HCl, pH 8.0, 150 mM NaCl, 1 mM EDTA) with 0.5 mL/min flow. The column was washed with strep wash buffer until a stable baseline was reached.

The protein was eluted with a step gradient to 100% strep elution buffer (100 mM Tris-HCl, pH 8.0, 150 mM NaCl, 1 mM EDTA, 2.5 mM desthiobiotin). 5 mL of the purified pG was injected in Superdex S75 16/60 equilibrated in SEC buffer (100 mM Tris-HCl, pH 8.0, 150 mM NaCl, 1 mM EDTA, 2 mM TCEP) with a flow rate of 1 mL/min. After size exclusion chromatography, fractions were pooled. The pooled fractions had a concentration of 2.2 mg/mL determined with spectrophotometry. Additionally, protein molecular weight and concentration were measured by SEC-MALS and UV-Vis spectrophotometry. Purified pG was diluted to 50 µM in 350 mM NaCl and flash frozen in liquid nitrogen and stored at −80 °C.

### pG - oligo (non-cnvK) synthesis

The coupling of pG to a non-cnvK modified oligonucleotide barcode was performed following a previously described protocol^38^ with modifications. 10 nmol solution of 5′-aminated barcode oligonucleotide (IDT, resuspended from desalted powder into 1 mM stock) was reacted with 100 nmol freshly dissolved sulfo-SMCC (10 mg/mL in anhydrous dimethylformamide, freshly prepared from Pierce No-Weight format, Thermo, A39268) in PBS (pH 7.2, Thermo, 28372, dissolved from the powder in nuclease-free water) in DNA LoBind tube. The reaction (10 µL oligo, 85 µL PBS, 5 µL sulfo-SMCC) was incubated for 2.5 h at room temperature, 40 rpm, protected from light. The reaction was quenched with 1 µL 1 M Tris-HCl (pH 7.0). The reaction mixture (100 µL) was supplemented with glycogen (3 µL, 5 µg/µL), NaCl (10 µL, 5 M), and ethanol (300 µL, −20 °C). Samples were incubated at −70 °C for 30 min, centrifuged (19,000 × g, 30 min, 4 °C), resuspended in PBS (100 µL), and precipitated again with NaCl and ethanol. Pellets were washed in 90% ethanol (800 µL), centrifuged (19,000 × g, 15 min, 4 °C), and air-dried (5 min). pG buffer was exchanged on 7 MWCO Zeba Spin Desalting Columns to remove reducing agents. Columns were equilibrated with 4 x 350 µL PBS (pH 7.2) washes (centrifuge 1,000 x g, 2 min). 100 µL of pG (50 µM solution in 100 mM Tris/HCl pH 8.0 supplemented with 150 mM NaCl, 1 mM EDTA, 2 mM TCEP) was loaded onto the column and eluted by centrifugation (1,000 x g, 2 min) into a new protein low-binding tube. Desalted pG was used immediately. The air-dried oligonucleotide pellet was reconstituted in 74.8 µl of 1x PBS (pH 7.2) and 25.2 µl (1 nmol) of desalted Prot G (final molar ratio is 10:1 oligo: pG), protected from light. The reaction was incubated overnight at 4 °C, 22 rpm followed by quenching with 1.5 µl of 10 mM DTT (no Weight Format, Thermo, A39255) for 15 min, room temperature, 25 rpm. DTT was removed by 7K MWCO Zeba Spin Desalting columns, as was described previously. The resulting pG-oligo conjugate was immediately used for antibody conjugation or flash-frozen in liquid nitrogen and stored at -70 °C.

### UnO oligonucleotide design

The universal adapter oligo was designed according to the following criteria: (1) efficient hybridization at room temperature (length chosen to be 14 nucleotide), (2) low sequence similarity to commonly used published barcode sequences (Immuno-SABER, CODEX, TotalSeq - not more than 4-mer overlap), (3) balanced GC-content (40-60 %), (4) flanking G/C to enable stable binding, (5) lack of secondary structure, and finally (6) lack of homopolymer regions (not more than 3 nucleotides in a row). EGNAS^66^ was used to design oligonucleotide satisfying (1)-(6), yielding two sequences satisfying all of the criteria: 5’-GCATCCA<u>G</u>TCTCAG-3’ and 5’-GGAAGAACAGCAAG-3’. The first sequence was arbitrarily selected as a universal barcoding oligo *U* (<u>underlined</u> - cnvK position); second sequence was reserved for optional pull-down purification of the conjugated antibody^6^*. U* oligo (final sequence: 5’-GCATCCA[cnvk]TCTCAG-3’) was ordered at 2 µmol synthesis scale with reverse-phase chromatography purification.

### UnO synthesis

For UnO synthesis, cnvK-modified universal adapter oligo (*U*-oligo) (ordered from Metabion or Genelink), resuspended from desalted powder into 1 mM stock in nuclease-free water) was functionalized with sulfo-SMCC, and pG was desalted as described in “pG - oligo conjugation” section. 10 nmol of functionalized *U*-oligo was reacted with 25 nmol of the desalted pG (final molar excess is 2.5:1 pG : cnvK oligo) overnight.

### UnO purification

400 µL of Pierce Streptavidin Magnetic Beads (10 mg/mL) were placed into 2 mL DNA LoBind tubes and washed 3x in 1 mL of modified Binding/Wash (BW) buffer (20 mM Tris, 50 mM KCl, 0.1% Tween-20, nuclease-free water;) on the magnetic stand (Thermo DynaMag-2 Magnet, 12321D). 20 µL of biotinylated capture oligonucleotide (1 mM) was added to 400 µL beads in the BW buffer and incubated overnight at 4 °C with rotation (22 rpm). Beads were equilibrated to room temperature with rotation (22 rpm) and blocked by addition of 3 µL of nuclease-free BSA (50 mg/mL, RNase-free, Thermo, AM2616) for 10 minutes at room temperature with rotation (22 rpm), followed by washing 3 times in the BW buffer. After final wash beads were resuspended in 500 µL of BW buffer.

100 µL of unpurified UnO was added to 500 µL of the beads and incubated at 37 °C with shaking for 2 min. The tube was then placed on ice for 10 min, followed by 30 min incubation at 4 °C with rotation (22 rpm). Tubes were placed onto a pre-cooled magnetic stand (4 °C), incubated for 5 min to collect the beads, and supernatant (flow-through) was kept for SDS-PAGE analysis. Beads were washed 3x in 1 mL of BW buffer. In preparation for the elution, after the last wash beads are resuspended in 400 µL of new BW buffer, and the magnetic stand is heated to 37 °C in the incubator for 10 min. For the elution, the bead suspension is then transferred to 37 °C with shaking for 3 min and then placed into the pre-warmed magnetic rack to collect the supernatant containing purified UnO. Purified UnO was immediately used for antibody conjugation or flash-frozen in liquid nitrogen and stored at -70 °C.

### mUnO synthesis

mUnO (mIgG1-targeting UnO) comprised of (AlphaThera mIgG1 azide-substituted oYo-Link and *U*-oligo) was synthesized using copper-free click coupling of a DBCO-functionalized cnvK oligonucleotide to azide-modified oYo-Link mIgG1 binder (AlphaThera). To incorporate DBCO group, 5 µL of 5′-aminated *U-oligo* with cnvK (1 mM stock in nuclease-free water) was diluted in 93 µL of bicarbonate conjugation buffer (150 mM NaHCO₃, pH 8.0; Thermo, J62495.AP) and reacted with 2 µL of 10-fold molar excess of 25 mM EZ-Link™ TFP Ester-PEG4-DBCO (Thermo, C20043), freshly dissolved to 25 mM in anhydrous DMSO. The reaction was incubated for 2.5 h at room temperature (23 °C), 40 rpm, protected from light. The reaction was quenched with the addition of 5 µL of 1 M Tris-HCl (pH 8.0) for 15 min at room temperature.

DBCO-functionalized U-oligo with cnvK was purified by ethanol precipitation. 10 µL of 5M NaCl (final 0.3 M) and 300 µL of ice-cold ethanol (3x reaction volume) were added to the reaction mix, and samples were incubated at −70 °C for 15 min, followed by centrifugation (19,000 x g, 30 min, 4 °C). Pellets were resuspended in PBS (100 µL) and precipitated a second time under identical conditions. Pellets were washed twice with 90% ethanol, centrifuged (19,000 x g, 15 min, 4 °C), and air-dried for 5 min.

Azide-substituted mIgG1 oYo-Link (lyophilized powder, lot 23058, AlphaThera) was reconstituted in 100 µL to 33 uM concentration in nuclease-free water according to manufacturer’s recommendations (equivalent to the amount sufficient to conjugate 100 µg of the antibody). Purified DBCO-functionalized cnvK oligonucleotide pellet was reconstituted directly in the freshly prepared oYo-link solution (oligo-DBCO : mIgG1 oYo-Link molar ratio 4:1, recommended range 1.5:1 to 10:1). The click reaction was incubated overnight at 4 °C with rotation (22 rpm).

Excess unconjugated oligonucleotide was removed using Zeba™ Spin Desalting Columns (7 kDa MWCO), pre-equilibrated with PBS, as described previously (“pG - oligo conjugation”). The reaction mixture was applied to the column and centrifuged according to the manufacturer’s instructions (1,000 × g, 2 min, room temperature). The flow-through containing purified mUnO was collected in protein low-binding tubes and used immediately for antibody conjugation stored at 4 °C for short-term use or at −20 °C in the Storage Buffer for long-term storage.

### pG - oligo antibody conjugation with UV crosslinking

All steps were performed on ice and under minimal light exposure, unless stated otherwise. For conjugation, a UV crosslinker box (365 nm, 2.4 J; Carl Roth 1777.1) was used.

16 µL of 50 µM stock of pG coupled to oligonucleotide barcode (∼800 pmol) was thawed on ice, mixed with 5 µg of antibody (∼40 pmol) and adjusted to 50 uL with 1x PBS supplemented with 350 mM NaCl in PCR tube. The reaction mixture was incubated at 37 °C for 5 min in a thermocycler (lid and block heated). Tubes were then wrapped in foil and rotated at 22 rpm for 10 min at room temperature.

Tubes were placed upright on a pre-chilled metal rack on ice, placed in the crosslinker box with caps open and positioned adjacent (< 1 cm) to LED, and irradiated for 3 min. Then lids were closed, and samples were returned to the crosslinker in a horizontal orientation for an additional 3 min of irradiation through the tube walls. Thereby we tried to compensate for any potential negative effects that may come from the UV absorbance of the polypropylene tubes and uneven absorbance of the barcoding solution volume, although this 2-step irradiation may not be strictly needed. For negative “UV−” control, samples were incubated in dark for 6 min in total.

As alternative UV sources, a simple commercial UV box (like a nail polish fixer, MelodySusie Ultraviolet Radiation Lamp 36W, DR-301C; crosslinking for 1h on ice with caps closed) or UV gun (UV Curing LED System, 365 nm, Thorlabs, CS20K2; crosslinking for 90 seconds, directing the beam around the full circumference of the tube) can be used. We found the latter method to be the least practical due to the difficulty of achieving uniform illumination and when processing multiple tubes.

### UnO antibody conjugation and optional purification

All steps were performed on ice and under minimal light exposure. For UnO, 16 µL of 50 µM stock of pG coupled to oligonucleotide barcode (∼800 pmol) was thawed on ice, mixed with 5 µg of antibody (∼40 pmol) and adjusted to 50 uL with 1x PBS supplemented with 350 mM NaCl in PCR tube. For mUnO: 1 µl of 33 µM purified defrosted mUnO (∼33 pmol) was used for 1µg of mIgG1 antibody (∼6,6 pmol) stock, following the manufacturer’s instructions for mIgG1 oYo-Link. For UnO and mUnO, before adjusting the final reaction volume to 50 µL, 4 µL of 200 µM barcode oligonucleotide solution in 1x PBS (800 pmol) was added. The reaction mixture was incubated at 37 °C for 5 min in a thermocycler (lid and block heated). UV crosslinking was performed as described above. Optionally, 14-nucleotide “blocker” oligos complementary to the barcode sequence (but not overlapping with the cnvK-crosslinking sequence) could be added to the photocrosslinking solution (final concentration 1 µM) to make the barcode double-stranded, potentially reducing the non-specific binding the oligos to the antibody (particularly for longer barcodes). Such double-stranded barcodes do not impact downstream analysis with SDS-PAGE or Immuno-SABER.

Excess of the oligonucleotide and non-reacted pG were removed using Amicon Ultra-0.5 100 kDa centrifugal filters (Merck Millipore, UFC5100) pre-blocked with 500 µL of 1x PBS containing 0.1% Tween-20 and centrifuged at 12,000 × g for 2 min at room temperature. The conjugation reaction was loaded onto the filter, topped with 250 µL 1x PBS containing 1 M NaCl (final volume 450 µL), and centrifuged at 12,000 × g for 8 min. Filters were washed three times with 450 µL cold 1x PBS containing 1 M NaCl (12,000 × g, 8 min each) to remove free DNA, followed by a single wash with 450 µL cold 1x PBS to remove excess salt. Concentrated protein was recovered by inverting the filter into a clean collection tube and centrifuging at 2,000 × g for 2 min, yielding 20–50 µL final volume. An aliquot (4 µL) was used for SDS–PAGE analysis. The remaining sample was mixed with 4x Storage Buffer (4x PBS, 5% nuclease-free BSA, 0.2% sodium azide, 20 mM EDTA, 0.2% Triton-X100) and glycerol (volume ratio sample:4x Storage Buffer:glycerol 1:1:2) and kept at −20 °C in PCR (small volume) or protein LoBind tubes for long-term storage.

### DNA and protein PAGE

Purified, unpurified conjugated antibodies and unconjugated controls were analyzed by NuPAGE 3-8% tris–acetate (non-reducing conditions; Thermo, EA03752BOX) or 4-12% bis-tris (under reducing conditions; Thermo, NP0329BOX) mini gels. Total protein was visualized using Krypton Protein Stain (Thermo, 46630), and single-stranded DNA was detected using SYBR Gold (Thermo, S11494). Antibody samples (400 ng per lane) were diluted in 1x LDS sample buffer (Thermo, NP0007) to 8 µL total volume. Samples were incubated at 72 °C for 10 min, cooled on ice for 3 min, and centrifuged briefly. Gel duplicates were prepared to be stained with either Krypton (protein) or Sybr Gold (DNA). A 10 µL aliquot of mPAGE Prestained Protein Ladder (Merck MPSTD4) or PageRuler™ Unstained Broad Range Protein Ladder (Thermo, 26630) was thawed at room temperature, mixed gently, and loaded at 5 µL per lane. NuPAGE 1x Tris–acetate SDS running buffer (Thermo, LA0041) was prepared from a 20x stock (45 mL + 865 mL deionized ice-cold water). Wells were rinsed to remove residual glycerol before loading. Electrophoresis was performed at 90 V for 10 min (stacking), followed by 150 V for 45 min. The running buffer was reused as noted, and gel tanks were washed thoroughly after use. Following electrophoresis, gels were removed from cassettes using a gel knife. The well region was excised. Gels were transferred directly into the fixation buffer (40% ethanol, 10% acetic acid in water). Gels were fixed for 30 min at room temperature with gentle agitation, the solution was replaced, and fixation was continued for another 30 min. For protein staining, gels were incubated with 1x Krypton Protein Stain (diluted from 10x stock; 50 mL per gel) for 1 h at room temperature. Gels were destained twice in 7% acetic acid for 15 min each, followed by two 15 min washes in deionized water, and imaged on Typhoon FLA 9000 scanner using a 532 nm laser with LPG (575 LP) emission filter or iBright 1500 gel documenting system (Cy3/Alexa Fluor 555 channel). For DNA staining, SYBR Gold stock (10,000x) was diluted 1:10,000 in 1x TAE buffer to a total volume of 50 mL. Gels were incubated with the solution for 30 min at room temperature in the dark and imaged immediately (473 nm laser with LPB (510 LP) emission filter on Typhoon or Alexa Fluor 488/FITC channel).

### SDS-PAGE Gel quantification

To estimate the conjugation efficiency, protein bands on non-reducing gels were manually quantified using Gel Analyzer in Fiji (2.14.0/1.54f): the intensity of bands corresponding to conjugated antibodies were divided by sum of the intensities of conjugated and non-conjugated bands. Additionally, we provide a custom Fiji macro for automatic band detection and quantification as described below in the Github repository (see **Data and Code Availability**).

The script uses the Trainable Weka Segmentation plugin^67^ to segment the bands from the gel image. For this the user manually trains a classifier to distinguish background from bands. Following training, the classifier is applied to the input image and the resulting segmentation mask is binarized. Two iterations of morphological erosion are applied to reduce noise and small speckles. The size of the remaining objects is measured and a threshold of 1000 pixels is used to only include proper bands. For each band its area, intensity and centroid position are recorded and saved for further processing.

It is important to note that most protein stains, including Krypton, bind to proteins in gel through electrostatic interactions and hydrophobic patches, which might have less staining efficiency when proteins are modified with negatively charged DNA groups. This effect is expected to be less for the non-reduced antibody (and becomes even more significant for bands of smaller molecular weight, as in reducing gels) but may still lead to a slight underestimation of the conjugated antibody amount.

Alternatively, one could judge pG-based conjugation efficiency on reduced gels by calculating depletion of Heavy Chain /Light Chain ratio, since heavy and light chain are generated from the same complex and pG binds to the heavy chain exclusively. In this case depletion of non-conjugated HC can be calculated to estimate the conjugation efficiency.

### Mammalian cell culture

HeLa cells were grown in Dulbecco’s Modified Eagle Medium (DMEM; low glucose, 1 g/L) supplemented with 10% (v/v) fetal bovine serum (FBS), 1x GlutaMAX (2 mM L-alanyl-L-glutamine), and 1x penicillin–streptomycin (100 U/mL penicillin, 100 µg/mL streptomycin). Cells were grown at 37 °C, 5% CO2. For experiments 10,000 cells/well were seeded in 18-well LabTek chamber slides and grown overnight in the culture medium. Cells were washed with pre-warmed 1x PBS and fixed with room temperature 4% paraformaldehyde (PFA) in PBS for 30 min at room temperature. Fixed cells were quenched in 100 mM Tris-HCl (pH 8.0) for 10 min at RT and washed three times in PBS.

### Tissues

For CODEX experiments (**Supplementary Fig. 9**), microscopy slides with 2 µm FFPE human tonsil tissue sections were prepared by the Pathology Department at University Hospital Düsseldorf (Ethics approval 2025-3434 and 2024-3082). For other experiments, 5 µm human FFPE tonsil sections were sourced commercially from Amsbio.

### Chemically synthesized concatemeric amplifiers for SABER-Flex

Primary and Secondary concatemers (**Supplementary Table 2**) were ordered as ultramer DNA oligonucleotides (Integrated DNA Technologies) were ordered at a 4 nmol synthesis scale and supplied as dry pellets in individual tubes, then resuspended to 100 µM in IDTE buffer (pH 8.0) prior to use. These concatemeric probes can be used as synthesized for branched DNA amplification^53^ or can be optionally amplified prior to hybridization in order to increase the effective concentration and amount of the full-length probes using in vitro transcription mediated ssDNA amplification as was previously done for MERFISH^68^, incorporating further modifications that are previously described^69^. The detailed in vitro amplification protocol is provided below.

*Limited-Cycle PCR:* Limited-cycle PCR (LC-PCR) was performed using 1 µL of 100 µM ultramer stock as a template. A typical reaction (50 µL) contained 25 µL of 2x KAPA HiFi mix,10 µM forward and reverse primers (5 µL of each), and was amplified for 12 cycles. Cycling conditions were: 95°C 3 min; 12 cycles of 98°C 15 s, 54°C 15 s, 72°C 20 s; hold 4°C. Products were purified as above and assessed by agarose gel electrophoresis, Bioanalyzer, or TapeStation to confirm presence of a single band.

Forward primer: 5’GCC CCA TCA TGT GCC TTT CC

Reverse primer: T7-bringer REV GAA TTT AAT ACG ACT CAC TAT AGG GAG AGT GCC TGC GAG AGG GAA ATC CA

Both primers were ordered as 10 uM desalted oligos from IDT.

*In Vitro Transcription*: Amplified dsDNA templates containing a T7 promoter were transcribed using the HiScribe T7 kit (New England Biolabs). Typical reaction (20 µL) contained 800–900 of ng template DNA (up to 8 µL), 100 mM NTP mix (ATP, GTP, UTP, CTP; 2 µL each), 2 µL of 10x reaction buffer, and 2 µL of T7 RNA polymerase mix. Reactions were incubated overnight at 37°C with a heated lid (37°C).

Following transcription, residual template DNA was removed by addition of 80 µL of DNase I solution (2 units of DNAse I, 10 µL of DNAse I Buffer, 69 µL of water), followed by incubation for 20 min at 37°C in a thermocycler. RNA was purified using the Monarch RNA Cleanup Kit (NEB) according to manufacturer’s recommendations, eluted in 50 µL nuclease-free water, and quantified spectrophotometrically. Typical yields ranged from 300–400 µg RNA per 20 µL reaction.

Reverse Transcription and RNA Hydrolysis: Single-stranded DNA probes were generated by reverse transcription (RT). RNA (≤50 µg per 50 µL reaction; adjusted with nuclease-free water) was combined with 4 µL of RT primer (1 mM), 10 µL of 5x RT buffer, 2.5 µL of 10 mM dNTP mix (0.5 mM final concentration), 3 µL of RNase inhibitor (MBP-RNAsin, ∼40 U/µL, in-house produced), 3 µL of betaine (5 M stock), and 2.5 µL of reverse transcriptase (Maxima H-, Thermo). Reverse transcription reactions were performed in a thermocycler with a heated lid (105 °C) using the following program: 50 °C for 3 min, followed by 42 °C for 90 min; followed by 10 cycles of 50 °C for 2 min and 42 °C for 2 min; the reaction was terminated by incubation at 85 °C for 5 min, and samples were held at 4 °C until further processing. RNA templates were degraded by alkaline hydrolysis by addition of 6 µL of 0.5 M EDTA and µL of 1 M NaOH and incubation for 15 min at 90°C), followed by neutralization with 6 µL of 1 M HCl. THe reaction was purified using Monarch RNA Cleanup columns. ssDNA was eluted in 50 µL water, yielding ∼60 µg per two combined RT reactions. ssDNA was analyzed using NanoDrop 2000 and pre-cast 1% Agarose E-gel (Thermo, A42135).

### Primer exchange reaction (PER) for Immuno-SABER

PER was performed as previously described^6^. Reaction conditions are listed in **Supplementary Table 4**. PER products were purified on Qiagen MinElute PCR purification kit according to manufacturer’s protocol and eluted in 20 µL of nuclease-free water. Concatemer concentration was measured using NanoDrop (typically ∼100–500 ng/µL). Purified PER products were stored at -20 °C until use.

### Immuno-SABER/IF staining

Immuno-SABER was performed as described in a recently published improved protocol^70^. Fixed cells were blocked following the Immuno-SABER protocol using Blocking Buffer (1x PBS (pH 7.4), 2% nuclease- and protease-free BSA (Sigma, 126609-10GM), 0.1% dextran sulfate (Millipore, S4030), 0.5 mg/mL sheared salmon sperm DNA (Invitrogen, AM9680), 4 mM EDTA (Invitrogen, AM9260G), 0.8 µM mixture of blocking 42-nt oligonucleotides (bc42_79-83, as listed in^70^), and 0.1% Triton X-100 (Sigma, T8787)). Blocking Buffer (50 µL per well) was applied onto the sample for 3 x 10 min at room temperature. For all experiments, antibody dilution recommended by the vendor was used (assuming no losses during conjugation procedure, **Supplementary Table 5**). Antibody staining solution was prepared by diluting the conjugated antibodies in Blocking Buffer supplemented with barcode blocking oligonucleotides (14-nt sequences complementary to antibody barcodes) at a final concentration of 5 µM each. The solution was pre-incubated at 37 °C for 5 min in a PCR block and cooled down at room temperature for 2 minutes. The staining solution (50 µL per well) was then applied onto samples for 60 min at room temperature in a humidified chamber. Excess antibody was removed by washing for 10 min in Blocking Buffer, followed by three washes for 5 min each in PBS. For colocalization experiments, samples were additionally stained with conventional Cy3 or Alexa Fluor 647 coupled anti-rabbit IgG (H+L) polyclonal secondary antibodies (Jackson ImmunoResearch, cat. 711-165-152 and cat. 711-605152) diluted in Blocking Buffer (1:500). A complete list of oligonucleotides is provided in the **Supplementary Tables 2** (for non-cnvK pG), **3; antibody information is provided in Supplementary Tables 5** (subcellular markers for cultured cells), and **6** (cell type markers for tonsil staining).

Antibody-stained samples were post-fixed with 100 µL of 100 mM BS3 (Thermo, 21580) in PBS for 20 min at RT, washed in PBS (5 min), quenched with 100 mM NH_4_Cl in PBS for 5 min, and washed twice in PBS (5 min each). Samples were stored overnight at 4 °C if necessary. A 60% formamide solution (in 1x PBS) was freshly prepared and equilibrated to RT. Samples were washed three times for 5 min each in 1x PBS supplemented with 60% formamide to elute barcode blocking oligonucleotides, then washed twice for 5 min each in PBS to remove residual formamide.

For Immuno-SABER concatemer hybridization, hybridization buffer (2x SSC, 20% formamide, 10% dextran sulfate, 0.1% Tween 20, 0.5 mg/mL shared salmon sperm DNA, nuclease-free water). Samples were pre-incubated in 100 µL/well buffer for 10 min at 37 °C in the flat-top thermocycler with heated lid. In the meantime, hybridization buffer containing 20 ng/µL (corresponds to ∼66 nM) of each of the concatemers was prepared and incubated at 37 °C until use. Pre-heated concatemer mix was applied to the samples (50 µL/well) and incubated for 1 h at 37 °C in the flat-top thermocycler with heated lid (primary hybridization step). After incubation, samples were immediately washed with a pre-warmed 2x SSC buffer supplemented with 0.1% Tween 20, followed by wash with 45% formamide in PBS with 0.1% Tween 20, followed by three washes with 1x PBS with 0.1% Tween-20 for 10 min at 37 °C.

Imager hybridization solution (400 nM fluorophore-conjugated imager oligonucleotide in 1x PBS with 0.1% Tween-20) was applied to the sample at room temperature and incubated in dark for 30 min, followed by three washes with 1x PBS with 0.1% Tween-20 for 5 min at 37 °C (imager hybridization step). Samples were then stained with 1 µg/µL DAPI in 1x PBS with 0.1% Tween-20 for 5 min followed by two brief washes with the same buffer. For multiplexed experiments, samples were processed in four sequential staining-imaging-stripping cycles. In each cycle, samples were stained with imager oligonucleotides corresponding only to the targets revealed in that cycle (see **Supplementary Tables 4-5**). To minimize imager crosstalk, decoy imagers (non-fluorescent) with sequences corresponding to targets not revealed in the given cycle were included (as implemented previously^70^). For imager stripping in between cycles. 50% formamide in 1x PBS was applied for 5 min three times at RT Imagers were reapplied as above and image acquisition was performed immediately after imager probe hybridization.

### SABER-Flex concatemer and imager hybridization

Samples were prepared as described for Immuno-SABER until the concatemer hybridization step. Concatemer hybridization buffer (2x SSC, 20% formamide, 10% dextran sulfate, 0.1% Tween 20, 0.5 mg/mL shared salmon sperm DNA; 100 µL/well) was applied to samples and pre-incubated for 10 min at 37 °C in the flat-top thermocycler with heated lid. Primary amplifiers (directly after chemical synthesis or pre-amplified with in vitro transcription) were diluted to 150 nM each in the hybridization buffer, and the resulting probe solution was applied to the samples (50 µL/well) for 1 h at 37 °C in the flat-top thermocycler with heated lid (primary amplifier hybridization step), followed by two washes in washing buffer (45% formamide in PBS with 0.1% Tween 20) at room temperature (5 min per wash), followed by three washes with 1x PBS with 0.1% Tween-20 for 10 min at 37°C.

Following primary probes, secondary amplifier hybridization was performed. Secondary amplifier probes were diluted to 150 nM each in secondary concatemer hybridization buffer consisting of 2x SSC, 10% formamide, 0.1% Tween-20, 0.5 mg/mL sheared salmon sperm DNA, and 10% dextran sulfate, and the resulting probe solution was applied to the samples (50 µL/well). Samples were incubated for 1 h to overnight at 37 °C in the flat-top thermocycler with heated lid to allow hybridization of secondary concatemers to the primary concatemer scaffold (secondary concatemer hybridization step). Following hybridization, samples were washed twice in a washing buffer consisting of 15% formamide in 2x SSC at room temperature for 5 min per wash, followed by an additional wash in the same buffer at 37 °C for 15 min. The buffer then was exchanged to 1x PBS with 0.1% Tween-20.

For imaging, fluorescently labeled 15-mer readout imager oligonucleotides (**Supplementary Table 2**) were diluted to a final concentration of 1 µM each in 1x PBS supplemented with 0.1% Tween-20, and the resulting probe solution was applied to the samples (50 µL/well) for 30 min in a humidified chamber protected from light. Following hybridization, samples were washed once in washing buffer consisting of 1× PBS supplemented with 0.1% Tween-20 at 37 °C for 5 min, followed by two washes in 1× PBS at room temperature for 5 min per wash to remove excess imager oligonucleotides prior to imaging, followed by a staining with 1 μg/ml DAPI (Invitrogen, cat. #D1306) in 1xPBS for 10 min. Image acquisition was performed immediately after imager probe hybridization.

### Image acquisition for cultured cells

Cell samples were imaged in 1x PBS with 0.1% Tween-20 on Nikon Ti2 microscope equipped with Kinetix sCMOS camera and CrestOptics X-Light V3 spinning disk (50 µm pinhole size) using a 40X magnification air objective (CFI P-Apo 40x Lambda, numerical aperture 0.95). For excitation, Celesta light engine was used with the following laser lines: 405 nm—DAPI, 546 nm and 638 nm—IF and Immuno-SABER channels. 4 fields of view were arbitrarily chosen (and respective x,y coordinates recorded) for each well, the maximum intensity plane in the DAPI channel was identified as the central plane and a 10 µm z-stack (step 0.6 µm) around the central plane was acquired. Exchange rounds were done manually off the stage. The stage adaptor was not changed in between cycles to acquire the same regions of the well.

### Image analysis for cultured cells

Raw images were parsed with the BioIO (v.1.2.1) library. Image data was rechunked and saved as zarr arrays (v2.16.1). For the multicycle experiment, maximum intensity projected images were 2D-registered using DAPI channel as a reference with pyStackReg (v.0.2.7) library. For the combined IF/Immuno-SABER experiment, Pearson coefficient (PCC) was calculated on z-stacks across all imaging planes containing optical section of the cells (defined as 5 planes above and below maximum intensity plane, with z-step 0.6 µm). PCC was calculated on pixels within percentile range 50% - 99.99% to exclude background (i.e., outside cells) and outlier pixels, respectively. Seaborn (v0.13.2) “regplot” function was used to fit the linear regression model and scatterplot visualization. For visualization purposes, image channels with nuclear bodies (SP100, SC35, SON, coilin) 99-th percentiles were used as lower contrast values.

Pixel-level clustering of multiplexed images in **Supplementary Fig. S8** (maximum z-projections) was performed with self-organizing maps (SOM) using Spatiomic package^71^. To account image shift after registration, 10% of the image was removed from each border. A shared 50x50 self-organizing map was trained on 500,000 pixels per FOV. To enrich for rare pixels containing nuclear bodies (coilin, SP100, SON and SC35), 20% of the training set was sampled from high-intensity pixels (95^th^ – 99.999^th^ percentile) in the corresponding channels. The rest of the 80% training pixels were sampled randomly. Channel intensities were clipped (nuclear body markers – between 95th and 99.999^th^ percentile, the rest – between 25^th^ and 99.98^th^ percentile) and min-max normalized before training. SOM prototypes were grouped by Leiden clustering, and all retained image pixels were assigned through their nearest prototype. The background cluster (pixels outside of the cells) were identified by a lack of expression across all markers. Marker enrichment across clusters was assessed across all non-background pixels using GPU-accelerated Wilcoxon test in rapids-singlecell^72^.

### Image analysis for tonsil tissues (SABER-FLEX)

Analysis was performed in Fiji (2.14.0/1.54f). Raw z-stacks processed by median (radius 1.0 pixel) and gaussian (radius 1.0 pixel) filters, transformed to a single-plane images using maximum intensity projection, followed by conversion into composite multichannel RGB images.

### Sample preparation for CODEX

CODEX protocol was performed as described previously^73^, which was adapted from^28^, with modification of the oligonucleotide blocking solution BC4 in case of UnO-barcoded antibodies, as described below. Briefly, formalin-fixed paraffin-embedded (FFPE) tonsil sections (2 µm) were mounted on charged slides, baked, deparaffinized (xylene/ethanol series), rehydrated, and subjected to heat-induced epitope retrieval in Tris–EDTA (pH 9.0), followed by bleaching. Slides were equilibrated in staining buffer, blocked (Akoya blockers N/J/S plus BC4 oligo mix supplemented with additional oligonucleotide GCATCCAGTCTCAG complementary to u* on CODEX barcodes used for conjugation – see barcode sequences in **Supplementary Table 6**), and incubated overnight at 4 °C with a DNA-barcoded antibody cocktail (DNA barcodes were coupled using either UnO or as previously described as part of a CODEX protocol^28^. Post-staining fixation comprised sequential 1.6% PFA, cold methanol, and BS3 crosslinking, after which a flow cell was mounted onto each slide for subsequent imaging with Phenocycler Fusion (Akoya). Antibodies and oligonucleotides are listed in **Supplementary Table 6.**

### Image analysis for CODEX

The CODEX Processor software (Akoya) was used for raw data registration and stitching. Resulting qptiff files were parsed, preprocessed and visualized using spatialproteomics library (v0.6.1)^74^. For alignment of adjacent tissue sections, images were converted to SpatialData (v0.2.3) object^75^, visualized in napari (v0.5.2) and identical sets of landmarks were added as Points layers for each of the sections. Landmark-based affine registration (“spatialdata.transformations.Affine”) was applied to align adjacent sections in a common coordinate system. Representative regions-of-interest were cropped using spatialdata “query.polygon” function^75^. For marker positivity analysis, cells were segmented using DeepCell^76^, background was manually thresholded in each channel, and marker positivity was predicted using gating. To visualize the number of cells positive for each marker in adjacent sections, barplot was created using pandas.

## ACKNOWLEDGEMENTS

This work was supported by core funding from European Molecular Biology Laboratory and a research collaboration with GSK through the EMBL GenTechDev Platform. We thank Eileen Furlong, Rainer Pepperkok and Giovanna Bergamini for establishment of platform collaborations. We thank Glenn Cremers and Tom de Greef for sharing their protein G constructs and, EMBL Protein Expression and Purification Core Facility (Kim Remans and Jacob Scheurich) for the recombinant expression and purification. We thank Peng Yin, Ninning Liu and Jocelyn Kishi for early discussions about the approach and to Marc-Andrea Bärtsch and Dmytro Dvornikov for productive discussions about the tissue experiments. This work is in part supported by the Health + Life Science Alliance Heidelberg Mannheim and received state funds approved by the State Parliament of Baden-Württemberg (“MULTI-SPACE” and “Explore!Tech”).

## MATERIAL, DATA AND CODE AVAILABILITY

The data and code used for analysis are deposited on Zenodo and GitHub, respectively, and will be made publicly available upon publication. Plasmids encoding photoreactive protein G (originally provided by Tom de Greef Lab), as well as p-benzoyl-l-phenylalanine tRNA/tRNA synthetase, are available on Addgene (Addgene ID’s: 258623 and 31190, respectively).

## AUTHOR CONTRIBUTIONS

V.P. designed and performed experiments, prepared samples, generated and analyzed data, and drafted the manuscript.

Y.E. led and performed experiments, prepared samples, generated data, and contributed to manuscript drafting.

C.S., B.K., R.R. and T.M. contributed to experimental optimization and data generation.

F.S. developed the gel analysis plugin.

S.D. and P-M.B contributed to supervision.

S.K.S. developed the project concept, contributed to data interpretation, wrote the manuscript, and supervised the study.

All authors read and approved the manuscript.

## REFERENCES

1. Hickey, J.W., Neumann, E.K., Radtke, A.J., Camarillo, J.M., Beuschel, R.T., Albanese, A., McDonough, E., Hatler, J., Wiblin, A.E., Fisher, J., et al. (2022). Spatial mapping of protein composition and tissue organization: a primer for multiplexed antibody-based imaging. Nat. Methods 19, 284–295. 10.1038/s41592-021-01316-y.

2. Liu, Y., DiStasio, M., Su, G., Asashima, H., Enninful, A., Qin, X., Deng, Y., Nam, J., Gao, F., Bordignon, P., et al. (2023). High-plex protein and whole transcriptome co-mapping at cellular resolution with spatial CITE-seq. Nat. Biotechnol. 10.1038/s41587-023-01676-0.

3. Stoeckius, M., Hafemeister, C., Stephenson, W., Houck-Loomis, B., Chattopadhyay, P.K., Swerdlow, H., Satija, R., and Smibert, P. (2017). Simultaneous epitope and transcriptome measurement in single cells. Nat. Methods 14, 865–868. 10.1038/nmeth.4380.

4. Wang, Y., Woehrstein, J.B., Donoghue, N., Dai, M., Avendaño, M.S., Schackmann, R.C.J., Zoeller, J.J., Wang, S.S.H., Tillberg, P.W., Park, D., et al. (2017). Rapid Sequential in Situ Multiplexing with DNA Exchange Imaging in Neuronal Cells and Tissues. Nano Lett. 17, 6131–6139. 10.1021/acs.nanolett.7b02716.

5. Schürch, C.M., Bhate, S.S., Barlow, G.L., Phillips, D.J., Noti, L., Zlobec, I., Chu, P., Black, S., Demeter, J., McIlwain, D.R., et al. (2020). Coordinated Cellular Neighborhoods Orchestrate Antitumoral Immunity at the Colorectal Cancer Invasive Front. Cell 182, 1341–1359.e19. 10.1016/j.cell.2020.07.005.

6. Saka, S.K., Wang, Y., Kishi, J.Y., Zhu, A., Zeng, Y., Xie, W., Kirli, K., Yapp, C., Cicconet, M., Beliveau, B.J., et al. (2019). Immuno-SABER enables highly multiplexed and amplified protein imaging in tissues. Nat. Biotechnol. 37, 1080–1090. 10.1038/s41587-019-0207-y.

7. Goltsev, Y., Samusik, N., Kennedy-Darling, J., Bhate, S., Hale, M., Vazquez, G., Black, S., and Nolan, G.P. (2018). Deep Profiling of Mouse Splenic Architecture with CODEX Multiplexed Imaging. Cell 174, 968–981.e15. 10.1016/j.cell.2018.07.010.

8. Liu, Y., Yang, M., Deng, Y., Su, G., Enninful, A., Guo, C.C., Tebaldi, T., Zhang, D., Kim, D., Bai, Z., et al. (2020). High-Spatial-Resolution Multi-Omics Sequencing via Deterministic Barcoding in Tissue. Cell 183, 1665–1681.e18. 10.1016/j.cell.2020.10.026.

9. Walker, J.M., Kazempour Dehkordi, S., Fracassi, A., Vanschoiack, A., Pavenko, A., Taglialatela, G., Woltjer, R., Richardson, T.E., Zare, H., and Orr, M.E. (2022). Differential protein expression in the hippocampi of resilient individuals identified by digital spatial profiling. Acta Neuropathol. Commun. 10, 23. 10.1186/s40478-022-01324-9.

10. Niemeyer, C.M., Adler, M., and Wacker, R. (2005). Immuno-PCR: high sensitivity detection of proteins by nucleic acid amplification. Trends Biotechnol. 23, 208–216. 10.1016/j.tibtech.2005.02.006.

11. Choi, J., Love, K.R., Gong, Y., Gierahn, T.M., and Love, J.C. (2011). Immuno-hybridization chain reaction for enhancing detection of individual cytokine-secreting human peripheral mononuclear cells. Anal. Chem. 83, 6890–6895. 10.1021/ac2013916.

12. Horta, S., Neumann, F., Yeh, S.-H., Langseth, C.M., Kangro, K., Breukers, J., Madaboosi, N., Geukens, N., Vanhoorelbeke, K., Nilsson, M., et al. (2021). Evaluation of Immuno-Rolling Circle Amplification for Multiplex Detection and Profiling of Antigen-Specific Antibody Isotypes. Anal. Chem. 93, 6169–6177. 10.1021/acs.analchem.1c00172.

13. Gandin, V., Kim, J., Yang, L.-Z., Lian, Y., Kawase, T., Hu, A., Rokicki, K., Fleishman, G., Tillberg, P., Castrejon, A.A., et al. (2024). Deep-Tissue Spatial Omics: Imaging Whole-Embryo Transcriptomics and Subcellular Structures at High Spatial Resolution. bioRxiv, 2024.05.17.594641. 10.1101/2024.05.17.594641.

14. Hosogane, T., Casanova, R., and Bodenmiller, B. (2023). DNA-barcoded signal amplification for imaging mass cytometry enables sensitive and highly multiplexed tissue imaging. Nat. Methods 20, 1304–1309. 10.1038/s41592-023-01976-y.

15. di Michiel, M., Moch, H., Varga, Z., and Bodenmiller, B. (2023). Multielement Z-tag imaging by X-ray fluorescence microscopy for next-generation multiplex imaging. Nature.

16. Schueder, F., Lara-Gutiérrez, J., Beliveau, B.J., Saka, S.K., Sasaki, H.M., Woehrstein, J.B., Strauss, M.T., Grabmayr, H., Yin, P., and Jungmann, R. (2017). Multiplexed 3D super-resolution imaging of whole cells using spinning disk confocal microscopy and DNA-PAINT. Nat. Commun. 8, 2090. 10.1038/s41467-017-02028-8.

17. Unterauer, E.M., Shetab Boushehri, S., Jevdokimenko, K., Masullo, L.A., Ganji, M., Sograte-Idrissi, S., Kowalewski, R., Strauss, S., Reinhardt, S.C.M., Perovic, A., et al. (2024). Spatial proteomics in neurons at single-protein resolution. Cell 187, 1785–1800.e16. 10.1016/j.cell.2024.02.045.

18. Liu, N., Dai, M., Saka, S.K., and Yin, P. (2019). Super-resolution labelling with Action-PAINT. Nat. Chem. 11, 1001–1008. 10.1038/s41557-019-0325-7.

19. Schueder, F., Lara-Gutiérrez, J., Haas, D., Beckwith, K.S., Yin, P., Ellenberg, J., and Jungmann, R. (2021). Super-resolution spatial proximity detection with proximity-PAINT. Angew. Chem. Int. Ed Engl. 60, 716–720. 10.1002/anie.202009031.

20. Woo, S., Saka, S.K., Xuan, F., and Yin, P. (2024). Molecular robotic agents that survey molecular landscapes for information retrieval. Nat. Commun. 15, 3293. 10.1038/s41467-024-46978-2.

21. Vistain, L., Van Phan, H., Keisham, B., Jordi, C., Chen, M., Reddy, S.T., and Tay, S. (2022). Quantification of extracellular proteins, protein complexes and mRNAs in single cells by proximity sequencing. Nat. Methods 19, 1578–1589. 10.1038/s41592-022-01684-z.

22. Li, H., Ma, X., Shi, D., and Wang, P. (2025). Proximity ligation assay: From a foundational principle to a versatile platform for molecular and translational research. Biomolecules 15, 1468. 10.3390/biom15101468.

23. Schulte, S.J., Shin, B., Rothenberg, E.V., and Pierce, N.A. (2023). Multiplex, quantitative, high-resolution imaging of protein:protein complexes via hybridization chain reaction. bioRxiv, 2023.07.22.550181. 10.1101/2023.07.22.550181.

24. Jones, J.A., McMahon, N.P., Zheng, T., Eng, J., Chin, K., Kwon, S., Nederlof, M.A., Gray, J.W., and Gibbs, S.L. (2021). Oligonucleotide conjugated antibody strategies for cyclic immunostaining. Sci. Rep. 11, 23844. 10.1038/s41598-021-03135-9.

25. Agasti, S.S., Wang, Y., Schueder, F., Sukumar, A., Jungmann, R., and Yin, P. (2017). DNA-barcoded labeling probes for highly multiplexed Exchange-PAINT imaging. Chem. Sci. 8, 3080–3091. 10.1039/c6sc05420j.

26. Harada, A., Maehara, K., Handa, T., Arimura, Y., Nogami, J., Hayashi-Takanaka, Y., Shirahige, K., Kurumizaka, H., Kimura, H., and Ohkawa, Y. (2018). A chromatin integration labelling method enables epigenomic profiling with lower input. Nature Cell Biology 2018 21:2 *21*, 287–296. 10.1038/s41556-018-0248-3.

27. Wiener, J., Kokotek, D., Rosowski, S., Lickert, H., and Meier, M. (2020). Preparation of single- and double-oligonucleotide antibody conjugates and their application for protein analytics. Sci. Rep. 10, 1457. 10.1038/s41598-020-58238-6.

28. Black, S., Phillips, D., Hickey, J.W., Kennedy-Darling, J., Venkataraaman, V.G., Samusik, N., Goltsev, Y., Schürch, C.M., and Nolan, G.P. (2021). CODEX multiplexed tissue imaging with DNA-conjugated antibodies. Nat. Protoc. 16, 3802–3835. 10.1038/s41596-021-00556-8.

29. Caraccio, C., van de Klashorst, J., Cherkas, S., Ancel, S., Kempchen, T.N., Vazquez, G., Goltsev, Y., Wang, Y.X., Nolan, G.P., and Hickey, J.W. (2026). Comparative evaluation of antibody-oligonucleotide conjugation strategies for multiplexed imaging applications. Lab. Invest. 106, 104262. 10.1016/j.labinv.2025.104262.

30. Hadjabdelhafid-Parisien, A., Bitsch, S., Macarrón Palacios, A., Deweid, L., Kolmar, H., and Pelletier, J.N. (2022). Tag-free, specific conjugation of glycosylated IgG1 antibodies using microbial transglutaminase. RSC Adv. 12, 33510–33515. 10.1039/d2ra05630e.

31. Früh, S.M., Matti, U., Spycher, P.R., Rubini, M., Lickert, S., Schlichthaerle, T., Jungmann, R., Vogel, V., Ries, J., and Schoen, I. (2021). Site-specifically-labeled antibodies for super-resolution microscopy reveal in situ linkage errors. ACS Nano 15, 12161–12170. 10.1021/acsnano.1c03677.

32. Tumey, L.N. (2020). An overview of the current ADC discovery landscape. Methods Mol. Biol. 2078, 1–22. 10.1007/978-1-4939-9929-3_1.

33. Sograte-Idrissi, S., Schlichthaerle, T., Duque-Afonso, C.J., Alevra, M., Strauss, S., Moser, T., Jungmann, R., Rizzoli, S.O., and Opazo, F. (2020). Circumvention of common labelling artefacts using secondary nanobodies. Nanoscale. 10.1039/d0nr00227e.

34. Schueder, F., Rivera-Molina, F., Su, M., Kidd, P., Rothman, J.E., Toomre, D., and Bewersdorf, J. (2023). Unraveling cellular complexity with unlimited multiplexed super-resolution imaging. bioRxiv, 2023.05.17.541061. 10.1101/2023.05.17.541061.

35. Zhong, S., Wang, R., Gao, X., Guo, Q., Lin, R., and Luo, M. (2025). Modular DNA barcoding of nanobodies enables multiplexed in situ protein imaging and high-throughput biomolecule detection. Elife 14, RP105225. 10.7554/eLife.105225.

36. Schlichthaerle, T., Ganji, M., Auer, A., Kimbu Wade, O., and Jungmann, R. (2019). Bacterially derived antibody binders as small adapters for DNA-PAINT microscopy. Chembiochem 20, 1032–1038. 10.1002/cbic.201800743.

37. Jung, Y., Lee, J.M., Jung, H., and Chung, B.H. (2007). Self-directed and self-oriented immobilization of antibody by protein G-DNA conjugate. Anal. Chem. 79, 6534–6541. 10.1021/ac070484i.

38. Cremers, G.A.O., Rosier, B.J.H.M., Riera Brillas, R., Albertazzi, L., and de Greef, T.F.A. (2019). Efficient Small-Scale Conjugation of DNA to Primary Antibodies for Multiplexed Cellular Targeting. Bioconjug. Chem. 30, 2384–2392. 10.1021/acs.bioconjchem.9b00490.

39. Stiller, C., Aghelpasand, H., Frick, T., Westerlund, K., Ahmadian, A., and Karlström, A.E. (2019). Fast and Efficient Fc-Specific Photoaffinity Labeling To Produce Antibody-DNA Conjugates. Bioconjug. Chem. 30, 2790–2798. 10.1021/acs.bioconjchem.9b00548.

40. Hui, J.Z., Tamsen, S., Song, Y., and Tsourkas, A. (2015). LASIC: Light Activated Site-Specific Conjugation of Native IgGs. Bioconjug. Chem. 26, 1456–1460. 10.1021/acs.bioconjchem.5b00275.

41. Rosier, B.J.H.M., Cremers, G.A.O., Engelen, W., Merkx, M., Brunsveld, L., and de Greef, T.F.A. (2017). Incorporation of native antibodies and Fc-fusion proteins on DNA nanostructures via a modular conjugation strategy. Chem. Commun. 53, 7393–7396. 10.1039/c7cc04178k.

42. Niu, J., Hagen, J., Yu, F., Kalyuzhny, A.E., and Tsourkas, A. (2023). Labeling of phospho-specific antibodies with oYo-link® Epitope tags for multiplex immunostaining. Methods Mol. Biol. 2593, 113–126. 10.1007/978-1-0716-2811-9_7.

43. Yoshimura, Y., and Fujimoto, K. (2008). Ultrafast reversible photo-cross-linking reaction: toward in situ DNA manipulation. Org. Lett. 10, 3227–3230. 10.1021/ol801112j.

44. Schueder, F., Strauss, M.T., Hoerl, D., Schnitzbauer, J., Schlichthaerle, T., Strauss, S., Yin, P., Harz, H., Leonhardt, H., and Jungmann, R. (2017). Universal super-resolution multiplexing by DNA exchange. Angew. Chem. Int. Ed Engl. 56, 4052–4055. 10.1002/anie.201611729.

45. Jungmann, R., Avendaño, M.S., Woehrstein, J.B., Dai, M., Shih, W.M., and Yin, P. (2014). Multiplexed 3D cellular super-resolution imaging with DNA-PAINT and Exchange-PAINT. Nat. Methods 11, 313–318. 10.1038/nmeth.2835.

46. Guo, S.-M., Veneziano, R., Gordonov, S., Li, L., Danielson, E., Perez de Arce, K., Park, D., Kulesa, A.B., Wamhoff, E.-C., Blainey, P.C., et al. (2019). Multiplexed and high-throughput neuronal fluorescence imaging with diffusible probes. Nat. Commun. 10, 4377. 10.1038/s41467-019-12372-6.

47. Strotton, M., Hosogane, T., di Michiel, M., Moch, H., Varga, Z., and Bodenmiller, B. (2023). Multielement Z-tag imaging by X-ray fluorescence microscopy for next-generation multiplex imaging. Nat. Methods 20, 1310–1322. 10.1038/s41592-023-01977-x.

48. Takei, Y., Yun, J., Zheng, S., Ollikainen, N., Pierson, N., White, J., Shah, S., Thomassie, J., Suo, S., Eng, C.-H.L., et al. (2021). Integrated spatial genomics reveals global architecture of single nuclei. Nature 590, 344–350. 10.1038/s41586-020-03126-2.

49. Gandin, V., Kim, J., Yang, L.-Z., Lian, Y., Kawase, T., Hu, A., Rokicki, K., Fleishman, G., Tillberg, P., Castrejon, A.A., et al. (2025). Deep-tissue transcriptomics and subcellular imaging at high spatial resolution. Science 388, eadq2084. 10.1126/science.adq2084.

50. He, S., Bhatt, R., Brown, C., Brown, E.A., Buhr, D.L., Chantranuvatana, K., Danaher, P., Dunaway, D., Garrison, R.G., Geiss, G., et al. (2022). High-plex imaging of RNA and proteins at subcellular resolution in fixed tissue by spatial molecular imaging. Nat. Biotechnol. 40, 1794–1806. 10.1038/s41587-022-01483-z.

51. Manesse, M., Patel, K.K., Bobrow, M., and Downing, S.R. (2020). The InSituPlex® staining method for multiplexed immunofluorescence cell phenotyping and spatial profiling of tumor FFPE samples. Methods Mol. Biol. 2055, 585–592. 10.1007/978-1-4939-9773-2_26.

52. Kishi, J.Y., Schaus, T.E., Gopalkrishnan, N., Xuan, F., and Yin, P. (2018). Programmable autonomous synthesis of single-stranded DNA. Nat. Chem. 10, 155–164. 10.1038/nchem.2872.

53. Xia, C., Babcock, H.P., Moffitt, J.R., and Zhuang, X. (2019). Multiplexed detection of RNA using MERFISH and branched DNA amplification. Sci. Rep. 9, 7721. 10.1038/s41598-019-43943-8.

54. Kishi, J.Y., Liu, N., West, E.R., Sheng, K., Jordanides, J.J., Serrata, M., Cepko, C.L., Saka, S.K., and Yin, P. (2022). Light-Seq: light-directed in situ barcoding of biomolecules in fixed cells and tissues for spatially indexed sequencing. Nat. Methods 19, 1393–1402. 10.1038/s41592-022-01604-1.

55. Sauer-Eriksson, A.E., Kleywegt, G.J., Uhlén, M., and Jones, T.A. (1995). Crystal structure of the C2 fragment of streptococcal protein G in complex with the Fc domain of human IgG. Structure 3, 265–278. 10.1016/s0969-2126(01)00157-5.

56. Page, M., and Thorpe, R. (2002). Purification of IgG Using Protein A or Protein G. In The Protein Protocols Handbook, J. M. Walker, ed. (Humana Press), pp. 993–994. 10.1385/1-59259-169-8:993.

57. New England Biolabs Affinity of Protein A/G for IgG Types from Different Species. https://www.neb.com/en/tools-and-resources/selection-charts/affinity-of-protein-ag-for-igg-types-from-different-species.

58. Gut, G., Herrmann, M.D., and Pelkmans, L. (2018). Multiplexed protein maps link subcellular organization to cellular states. Science 361. 10.1126/science.aar7042.

59. Kennedy-Darling, J., Bhate, S.S., Hickey, J.W., Black, S., Barlow, G.L., Vazquez, G., Venkataraaman, V.G., Samusik, N., Goltsev, Y., Schürch, C.M., et al. (2021). Highly multiplexed tissue imaging using repeated oligonucleotide exchange reaction. Eur. J. Immunol. 51, 1262–1277. 10.1002/eji.202048891.

60. Steinek, C., Pachmayr, I., Strauss, S., Honsa, M., Kwon, J., and Jungmann, R. (2026). Dynamic Binder Exchange improves protein labeling efficiency in DNA-PAINT up to 15-fold. Angew. Chem. Int. Ed Engl., e18685. 10.1002/anie.202518685.

61. Navaratna, S., Hamblett, I., and Tonnesen, H.H. (2000). Photoreactivity of biologically active compounds. XVI. Formation and reactivity of free radicals in mefloquine. J. Photochem. Photobiol. B 56, 25–38. 10.1016/s1011-1344(00)00056-7.

62. Konrad, A., Karlström, A.E., and Hober, S. (2011). Covalent immunoglobulin labeling through a photoactivable synthetic Z domain. Bioconjug. Chem. 22, 2395–2403. 10.1021/bc200052h.

63. Yu, F., Järver, P., and Nygren, P.-Å. (2013). Tailor-making a protein a-derived domain for efficient site-specific photocoupling to Fc of mouse IgG₁. PLoS One 8, e56597. 10.1371/journal.pone.0056597.

64. Hui, J.Z., and Tsourkas, A. (2014). Optimization of photoactive protein Z for fast and efficient site-specific conjugation of native IgG. Bioconjug. Chem. 25, 1709–1719. 10.1021/bc500305v.

65. Park, J., Lee, Y., Ko, B.J., and Yoo, T.H. (2018). Peptide-DIrected Photo-cross-linking for site-specific conjugation of IgG. Bioconjug. Chem. 29, 3240–3244. 10.1021/acs.bioconjchem.8b00515.

66. Kick, A., Bönsch, M., and Mertig, M. (2012). EGNAS: an exhaustive DNA sequence design algorithm. BMC Bioinformatics 13, 138. 10.1186/1471-2105-13-138.

67. Arganda-Carreras, I., Kaynig, V., Rueden, C., Eliceiri, K.W., Schindelin, J., Cardona, A., and Sebastian Seung, H. (2017). Trainable Weka Segmentation: a machine learning tool for microscopy pixel classification. Bioinformatics 33, 2424–2426. 10.1093/bioinformatics/btx180.

68. Moffitt, J.R., Hao, J., Wang, G., Chen, K.H., Babcock, H.P., and Zhuang, X. (2016). High-throughput single-cell gene-expression profiling with multiplexed error-robust fluorescence in situ hybridization. Proc. Natl. Acad. Sci. U. S. A. 113, 11046–11051. 10.1073/pnas.1612826113.

69. Beckwith, K.S., Ødegård-Fougner, Ø., Morero, N.R., Barton, C., Schueder, F., Tang, W., Alexander, S., Peters, J.-M., Jungmann, R., Birney, E., et al. (2025). Nanoscale 3D DNA tracing in non-denatured cells resolves the Cohesin-dependent loop architecture of the genome in situ. Nat. Commun. 16, 6673. 10.1038/s41467-025-61689-y.

70. Reinhardt, R., Straka, T., Vierdag, W.-M., Jevdokimenko, K., Hecht, F., Pianfetti, E., Hudelmaier, T., Lai, H., Fouquet, W., Fahrbach, F., et al. (2026). Enabling high-plex spectral imaging via DNA-barcoded signal tuning and panel optimization. bioRxiv, 2026.03.18.709053. 10.64898/2026.03.18.709053.

71. Spatiomic https://spatiomic.org/latest/.

72. Dicks, S., Heumos, L., May, L., Jimenez, S., Angerer, P., Gold, I., Virshup, I., Fischer, F., Gill, M., Boerries, M., et al. (2026). GPU-accelerated single-cell analysis at scale with rapids-singlecell. arXiv [q-bio.GN]. 10.48550/arXiv.2603.02402.

73. Schniederjohann, C., Bruch, P.-M., Dietrich, S., and Neumann, F. (2025). Multiplexed immunophenotyping of lymphoma tissue samples. Methods Mol. Biol. 2865, 375–393. 10.1007/978-1-0716-4188-0_16.

74. Meyer-Bender, M., Voehringer, H., Schniederjohann, C., Koziel, S., Chung, E., Popova, E., Brobeil, A., Held, L.-M., Munir, A., Scverse Community, et al. (2025). Spatialproteomics - an interoperable toolbox for analyzing highly multiplexed fluorescence image data. bioRxiv. 10.1101/2025.04.29.651202.

75. Marconato, L., Palla, G., Yamauchi, K.A., Virshup, I., Heidari, E., Treis, T., Vierdag, W.-M., Toth, M., Stockhaus, S., Shrestha, R.B., et al. (2024). SpatialData: an open and universal data framework for spatial omics. Nat. Methods. 10.1038/s41592-024-02212-x.

76. Greenwald, N.F., Miller, G., Moen, E., Kong, A., Kagel, A., Fullaway, C.C., McIntosh, B.J., Leow, K., Schwartz, M.S., Dougherty, T., et al. (2021). Whole-cell segmentation of tissue images with human-level performance using large-scale data annotation and deep learning. Cold Spring Harbor Laboratory, 2021.03.01.431313. 10.1101/2021.03.01.431313.

