## Supplementary Information for "A dual-photoreactive adapter enables one-step DNA barcoding of antibody panels for multiplexed imaging"

**Supplementary Information** includes:

**Supplementary Figure S1.** Conjugation of rabbit monoclonal antibodies to protein G-based adapter coupled with DNA barcode oligos
**Supplementary Figure S2.** Multiplexed staining controls with SABER-Flex and conventional IF
**Supplementary Figure S3.** 3-plex SABER-Flex staining of 5 µm human FFPE tonsil section with barcoded rabbit monoclonal anti-CD3, CD8 and CD68 antibodies
**Supplementary Figure S4.** Micrograph of fixed HeLa cells stained with barcoded GM130, alpha-Tubulin and vimentin antibodies
**Supplementary Figure S5.** Supplementary Immuno-SABER colocalization experiments for a-Tubulin and GM130 antibodies
**Supplementary Figure S6.** SDS-PAGE analysis of UnO-conjugated antibodies
**Supplementary Figure S7.** Signal-to-background of individual markers in 10-plex Immuno-SABER experiment
**Supplementary Figure S8.** Unsupervised pixel clustering of the multiplexed images in Fig. 3D
**Supplementary Figure S9.** Application of UnO-barcoded antibodies for CODEX

**Supplementary Figure S10.** Non-reducing SDS–PAGE analysis of UnO-mediated barcoding of antibodies.

**Supplementary Table 1.** Overview of Immuno-SABER and SABER-Flex detection schemes
**Supplementary Table 2.** List of antibodies, protein G barcodes, SABER-Flex concatemers and the respective imagers.
**Supplementary Table 3.** List of Immuno-SABER barcode sequences.
**Supplementary Table 4.** List of Immuno-SABER primers and corresponding PER conditions.
**Supplementary Table 5.** List of Immuno-SABER antibodies, respective barcodes and primers.
**Supplementary Table 6.** List of CODEX antibodies and respective barcodes.

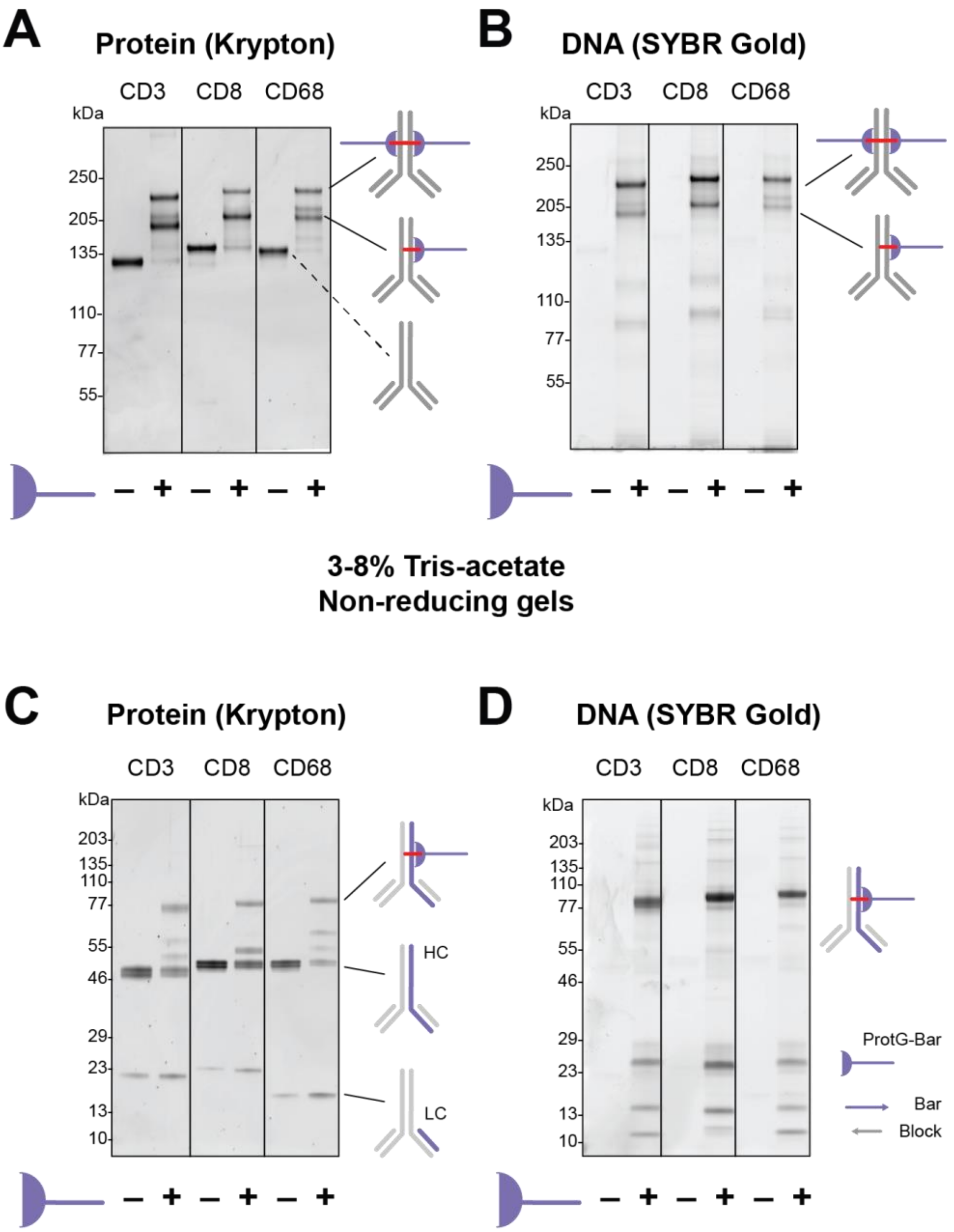


**Supplementary Figure S1. Conjugation of rabbit monoclonal antibodies to DNA barcodes through pG**. **(A-B)** Non-reducing 3-8% Tris-acetate PAGE gel for anti-CD3, CD8 and CD68 rabbit monoclonal antibodies stained with Krypton for protein visualization, and with SYBR Gold for DNA visualization. **(C-D)** Reducing 4-12% Bis-tris PAGE gel for the same antibodies allows seeing all heavy and light Chain products separately in more detail (while ProtG-barcode roughly amounts to ~17 kDa, and offers a visible shift in the non-reducing gel to evaluate the conjugation efficiency, the reducing gels are helpful to see the specific targeting of ProtG to Fc fragment of heavy chain, and the reaction subproducts). Schematics show the expected products (HC: heavy chain, LC: Light chain, pG-Bar: ProteinG conjugated with the barcoding oligo, Bar: barcode oligo, Block: optional 12mer ds-blocker oligo). Please note that Krypton staining of barcoded products might appear as weaker bands due to negative charge of DNA negatively affecting the Krypton stain, and this effect might be higher for reducing gels, causing underestimation of the conjugation efficiency. High conjugation efficiency is achieved for all three conjugates (on average 98, 96, 97%, respectively for CD3, CD8, CD68, n = 3 independent conjugations for each antibody).

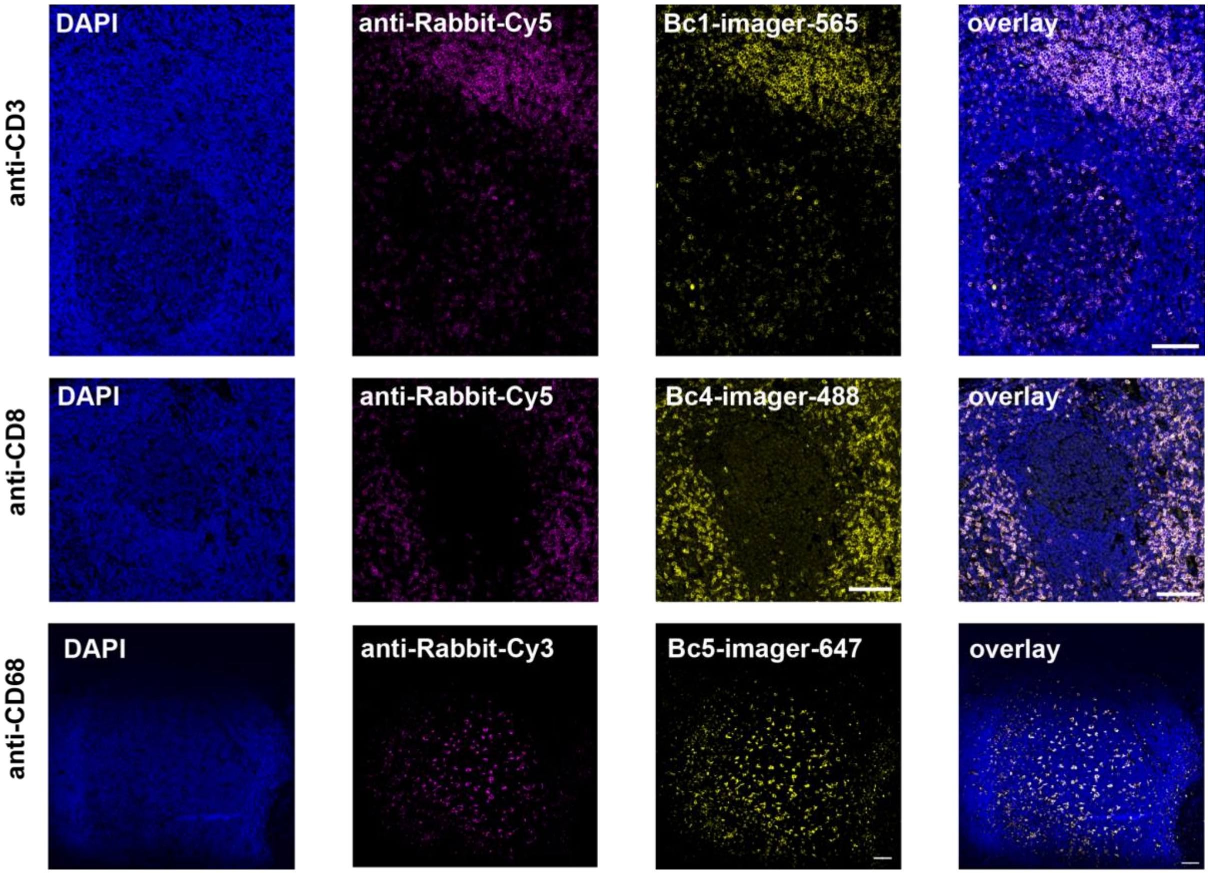


**Supplementary Figure S2. Multiplexed staining controls with SABER-Flex and conventional IF**. Combined SABER-Flex and conventional immunofluorescence staining with barcoded rabbit monoclonal anti-CD3, CD8 and CD68 antibodies in 5 µm human FFPE tonsil section. Immunostaining demonstrates similar staining patterns and expected distribution of positive cells (CD3+ and CD8+ T cells - outside of the germinal center, CD68+ macrophages - within the germinal center). Scale bar is 50 µm.

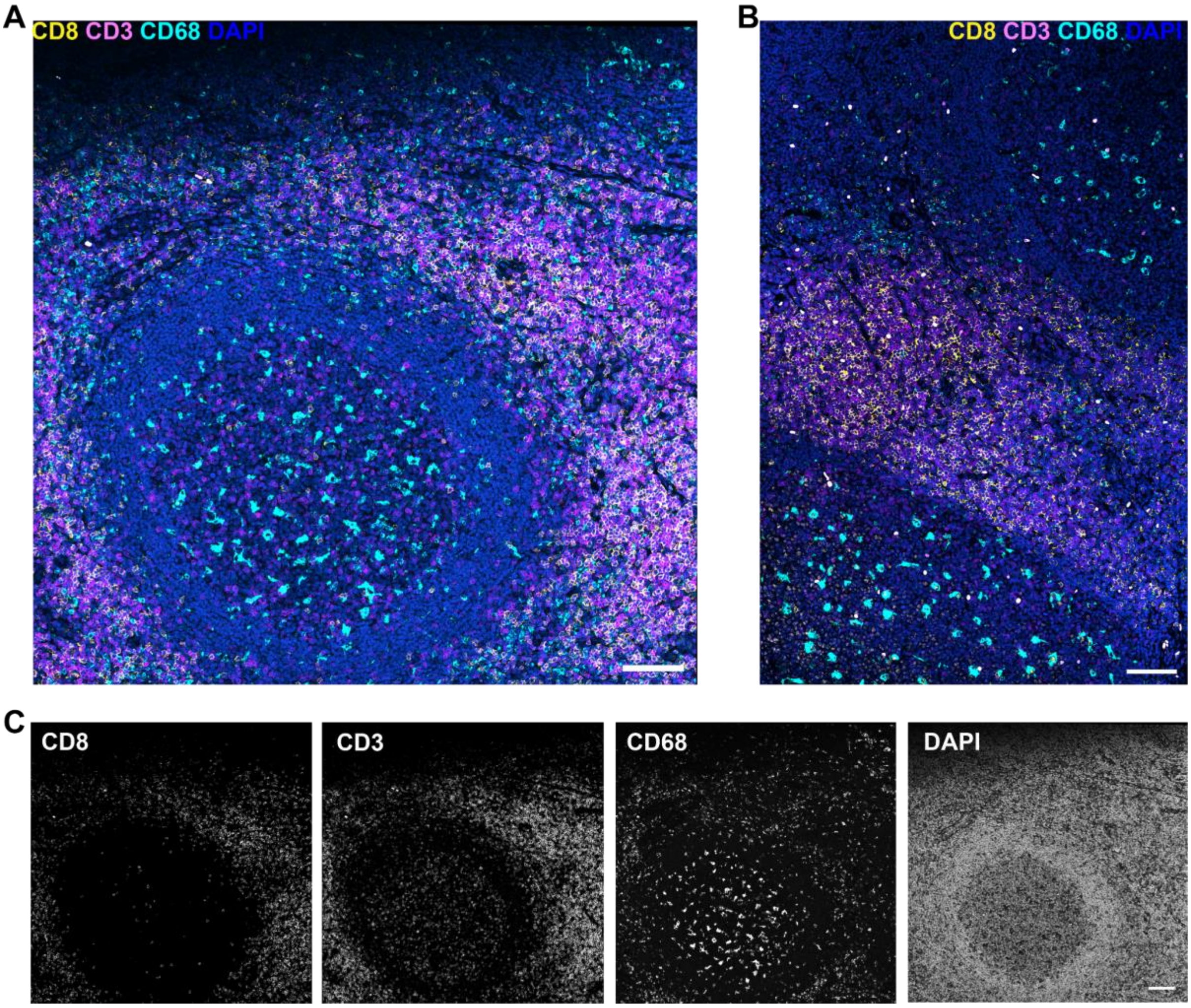


**Supplementary Figure S3. 3-plex SABER-Flex staining of 5 µm human FFPE tonsil section with barcoded rabbit monoclonal anti-CD3, CD8 and CD68 antibodies.** **(A, B)** Composite 4-color images of germinal centers (GC, in **A**) and extra-GC (in **B**) areas demonstrate distribution of T cells (CD3+, CD8+) and macrophages (CD68+). **(C)** Respective single-channel grayscale images corresponding to the field of view shown in (A). Scale bar is 100 µm.

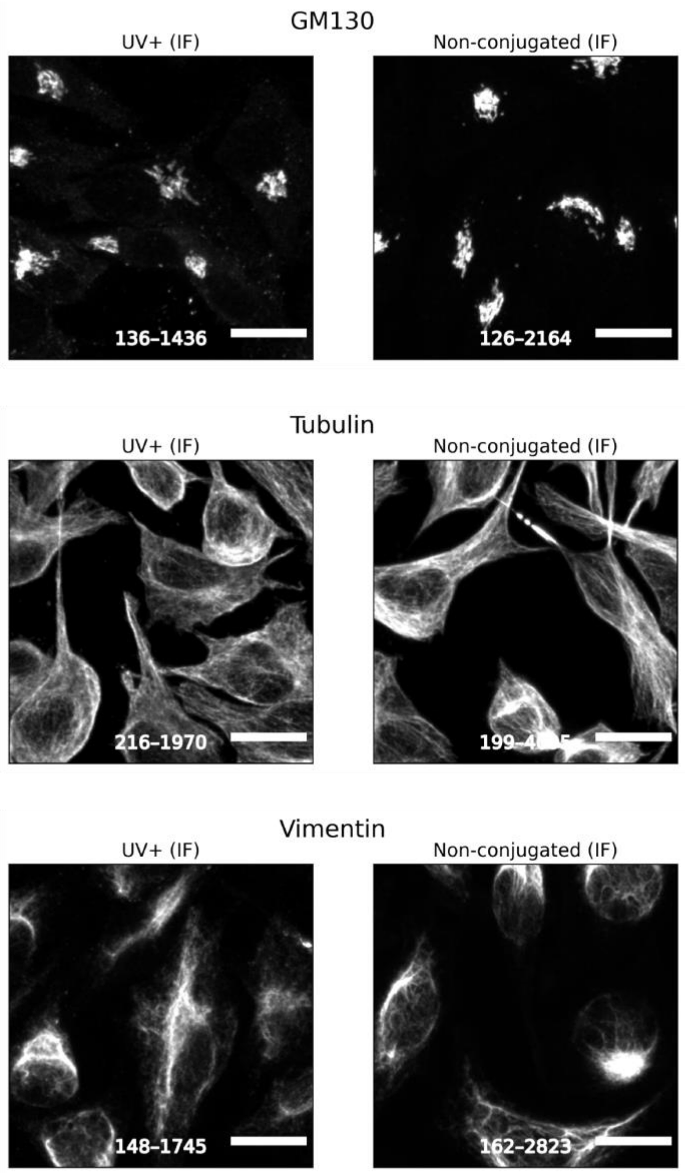


**Supplementary Figure S4**. **Micrograph of fixed HeLa cells stained with barcoded GM130, alpha-Tubulin and vimentin antibodies** ("UV+") or negative controls ("UV-" or "Non-conjugated"; see main text for reference). For "UV+" and "UV-" conditions both Immuno-SABER and conventional IF channels are shown. Intact primary antibodies ("Non-conjugated IF") demonstrate morphologically similar staining patterns as their UnO-barcoded counterparts and "UV-" controls. Images were auto contrasted based on bottom (5%) and top (99.9%) percentiles, contrast values are reported at the bottom of each image. Scale bar is 20 µm.

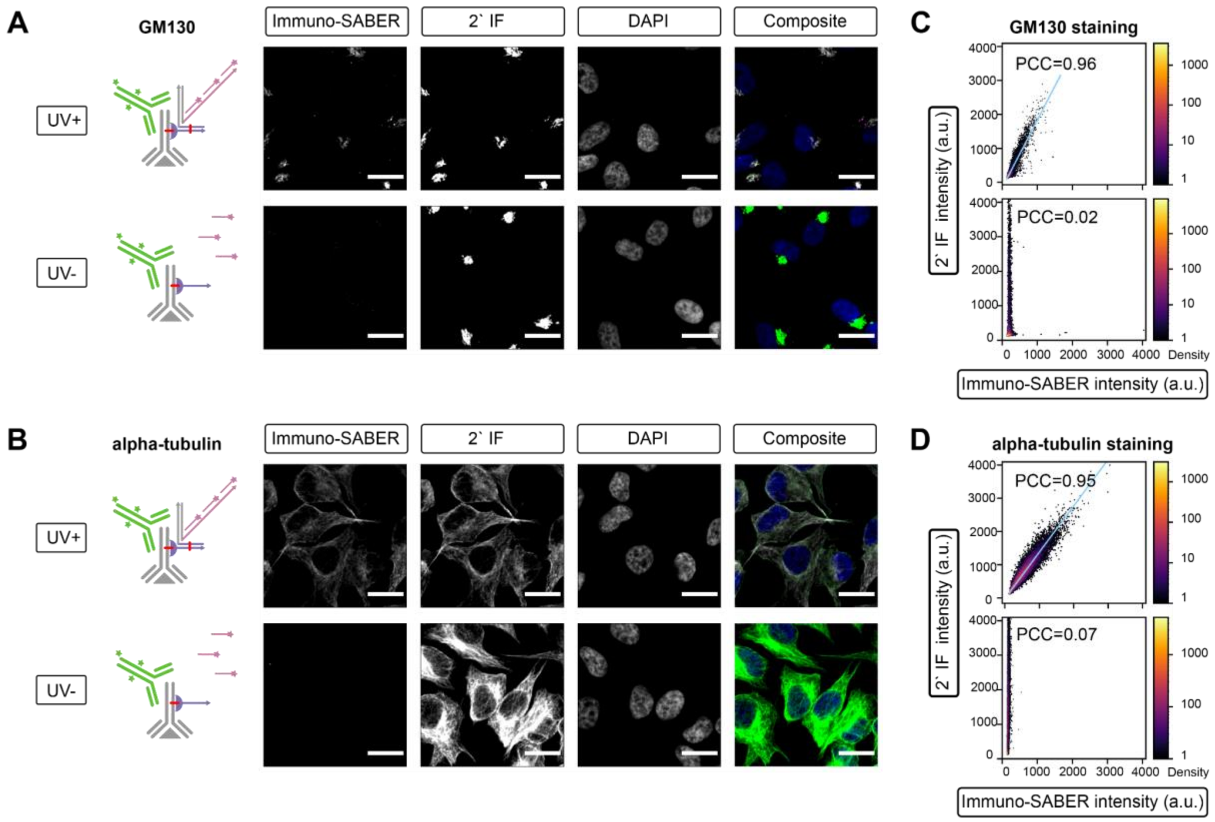


**Supplementary Figure S5. Supplementary Immuno-SABER colocalization experiments for alpha-Tubulin and GM130 antibodies**. **(A, B)** Fluorescent microphotographs of fixed HeLa cells stained with the respective primary antibodies (GM130 in **A**, alpha-Tubulin in **B**) conjugated with UnO and irradiated with UV ("UV+", top row) or non-irradiated control ("UV-", bottom row). Primary antibody staining is further revealed according to Immuno-SABER protocol with conventional indirect IF in parallel. Fluorescent channels correspond to (from left to right): Immuno-SABER (ATTO-565), indirect IF (Alexa Fluor 635 conjugated secondary antibody), DAPI (blue) and composite image (pink and green pseudo-color representing Immuno-SABER and IF channels, respectively, with white showing colocalization between two channels). Arrows on staining schematics correspond to 3' DNA end. **(C, D)** Respective fluorescence intensity scatterplots (raw intensity units) for Immuno-SABER (x-axis) and indirect IF (y-axis) channels for respective UnO "UV+" and "UV-" conditions. Pearson correlation coefficients are indicated on the plots. Blue line represents linear regression fit (*seaborn.regplot function*). Individual points are colored by local normalized density (log-scale color bar is indicated on the right). Scale bar is 20 μm.

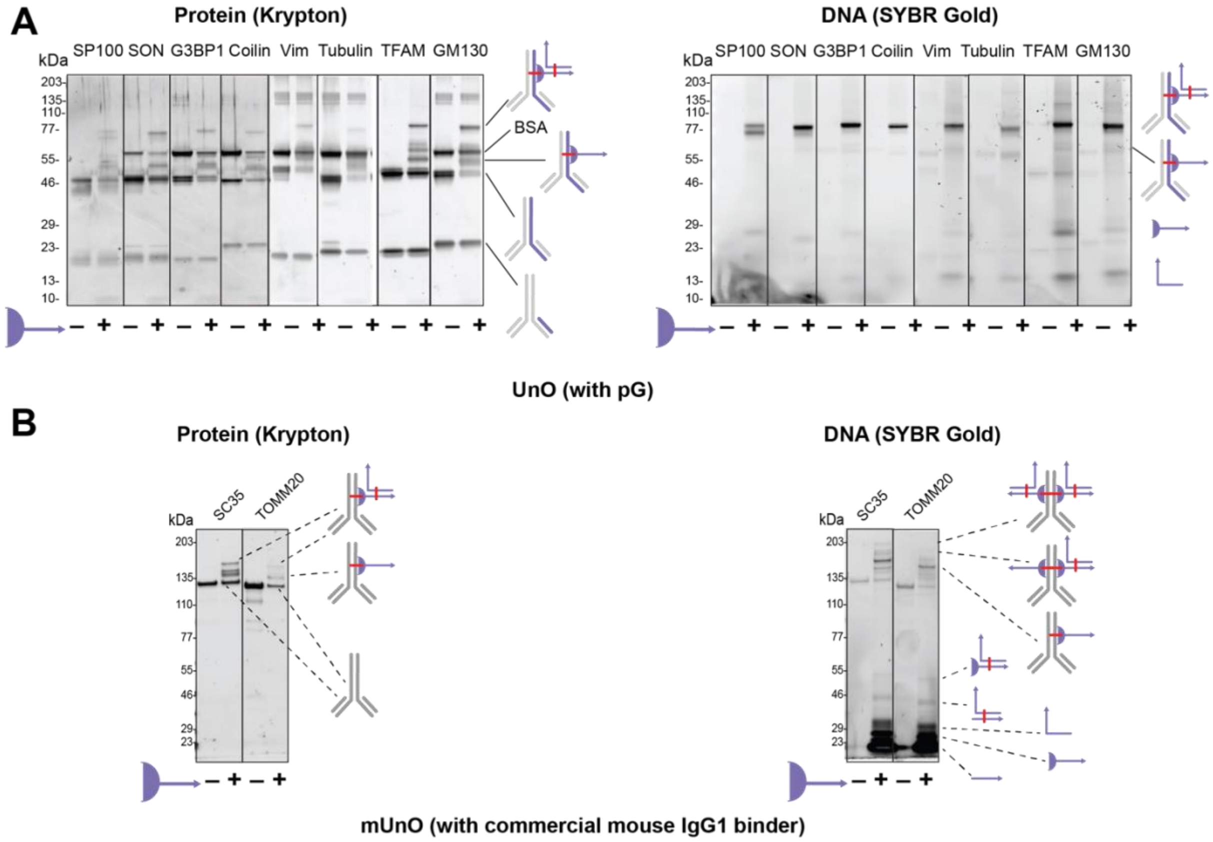


**Supplementary Figure S6. SDS-PAGE analysis of UnO-conjugated antibodies**. (**A**) Reducing SDS-PAGE (3-8% Tris-acetate gels) analysis of UnO-mediated barcoding of antibodies and the respective non-conjugated controls against (left to right): SP100, SON, G3BP1, Coilin, Vimentin, Tubulin, TFAM, and GM130. (**B**) Non-reducing (4-12% bis-tris) SDS-PAGE analysis of mUnO (mouse IgG1 oYo-link based) mediated barcoding of SC35 and TOMM20 and the respective non-conjugated controls. Gels were stained with krypton (protein, left) and SYBR Gold (DNA, right).

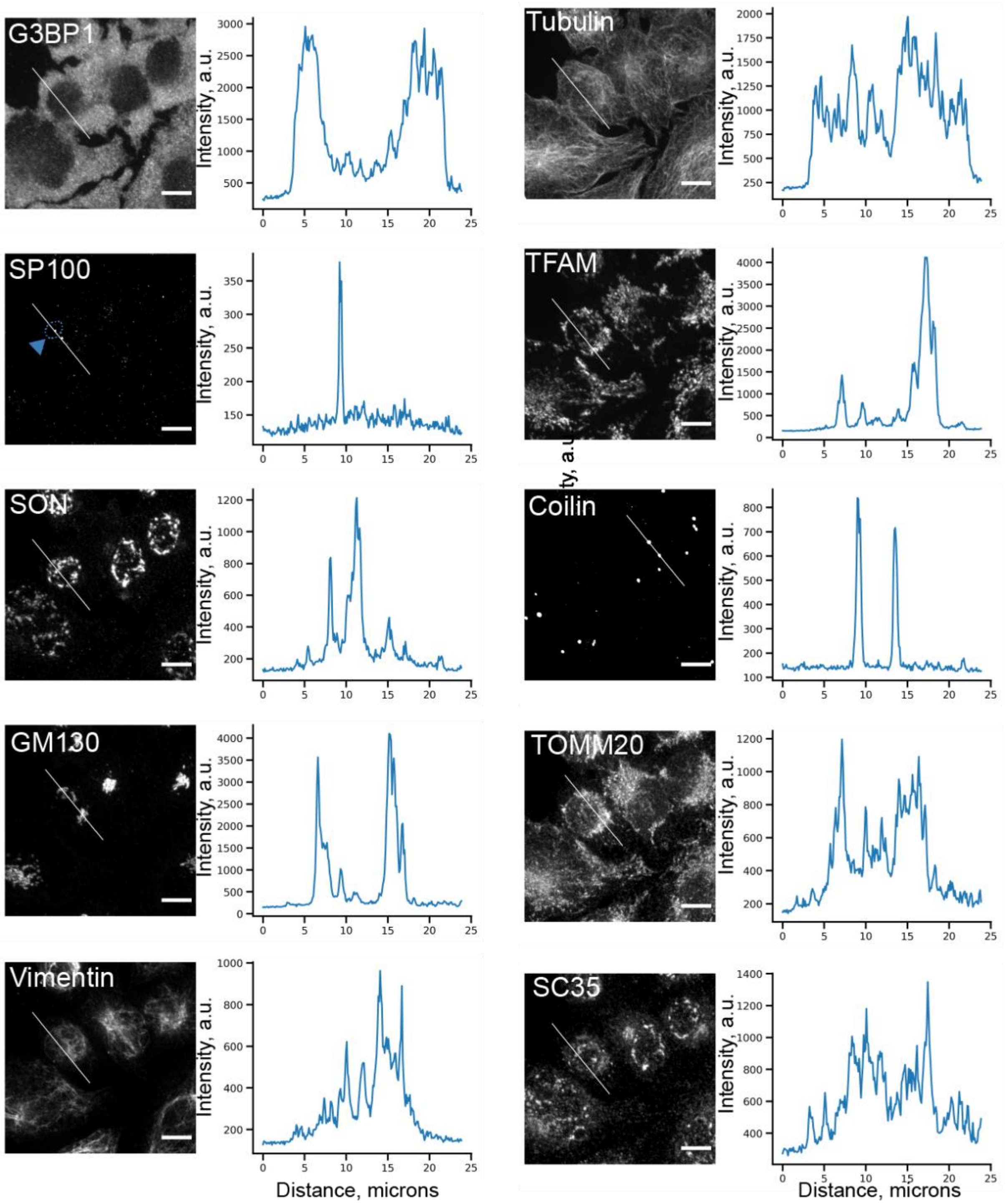


**Supplementary Figure S7. Signal-to-background of individual markers in 10-plex Immuno-SABER experiment**. Fluorescent images of fixed HeLa cells stained with: G3BP1, alpha-Tubulin, SP100, TFAM, SON, Coilin, GM130, TOMM20, Vimentin, SC35. Corresponding fluorescence intensity line scans showing the signal and background levels in arbitrary fluorescence units (a.u.) across the same 25 µm line (white) for all channels. The plots show well separated signal profiles corresponding to non-overlapping structures, for e.g. cytoplasmic G3BP1 versus nuclear SP100 or coilin. For SP100 fluorescent punctum intersected by the line scan is outlined with a blue dashed line and an arrowhead. Scale bar is 10 µm.

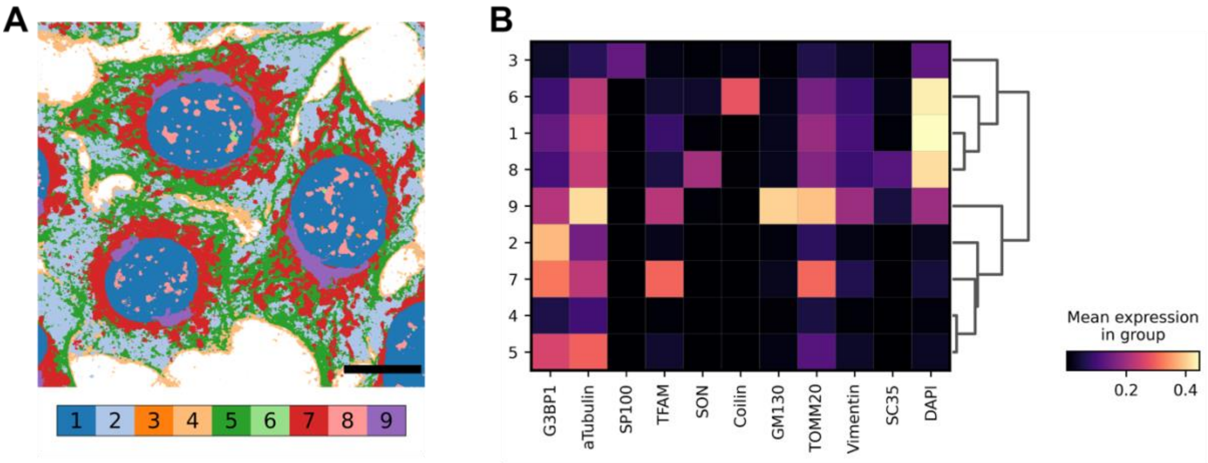


**Supplementary Figure S8. Unsupervised pixel clustering of the multiplexed images in Fig. 3D.** (**A**) the field of view from **Fig. 3D** with each pixel coloured by its assigned cluster, or multiplexed cell unit (MCU); the legend below lists MCUs and their assigned identities. Background pixels outside cells were identified by absence of signal across all markers and excluded. Scale bar: 20 µm. (**B**): mean marker intensity per MCU (rows) for each marker (columns), clipped and min-max normalized as described in the Methods. Clustering was performed without reference to marker identity and recovers compartments matching the known localizations of the panel such as mitochondria (MCU7; TFAM, TOMM20), Golgi (MCU9; GM130) and various nuclear bodies (speckles – MCU8; Cajal bodies – MCU6; PML bodies – MCU3). Because a diffraction-limited max-projected pixel may contain more than one structure, individual MCUs can receive contributions from several organelles. Analysis used 4 fields of view and ~24 Mio pixels; per-MCU marker enrichment was assessed by Wilcoxon test (**Methods**).

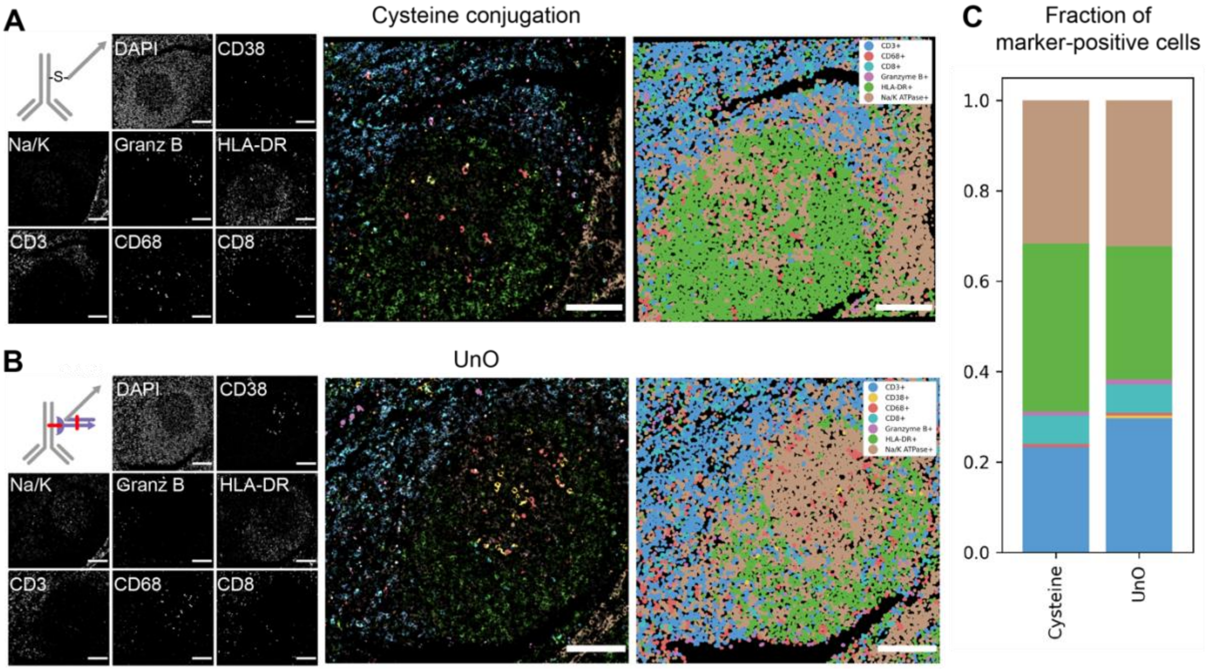


**Supplementary Figure S9. Application of UnO-barcoded antibodies for CODEX**. Antibodies against CD38, Na/K ATPase, Granzyme B, HLA-DR, CD68, CD8 and CD3 were conjugated according to cysteine-directed conjugation protocol **(A)** or UnO **(B)**. Both antibody panels were used for staining of adjacent tonsil sections and multiplexed imaging on PhenoCycler. Panels on the right represent cell masks, colored according to predicted positivity for each marker (see **Methods**). **(C)** Quantification of marker positivity for each cell in CODEX data. Barplot represents the fraction of cells positive for each marker for cysteine or UnO-conjugated antibodies. Note that custom carrier-free formulations were used for cysteine-directed conjugation and off-the-shelf antibodies were used for UnO. Scale bar: 100 μm.

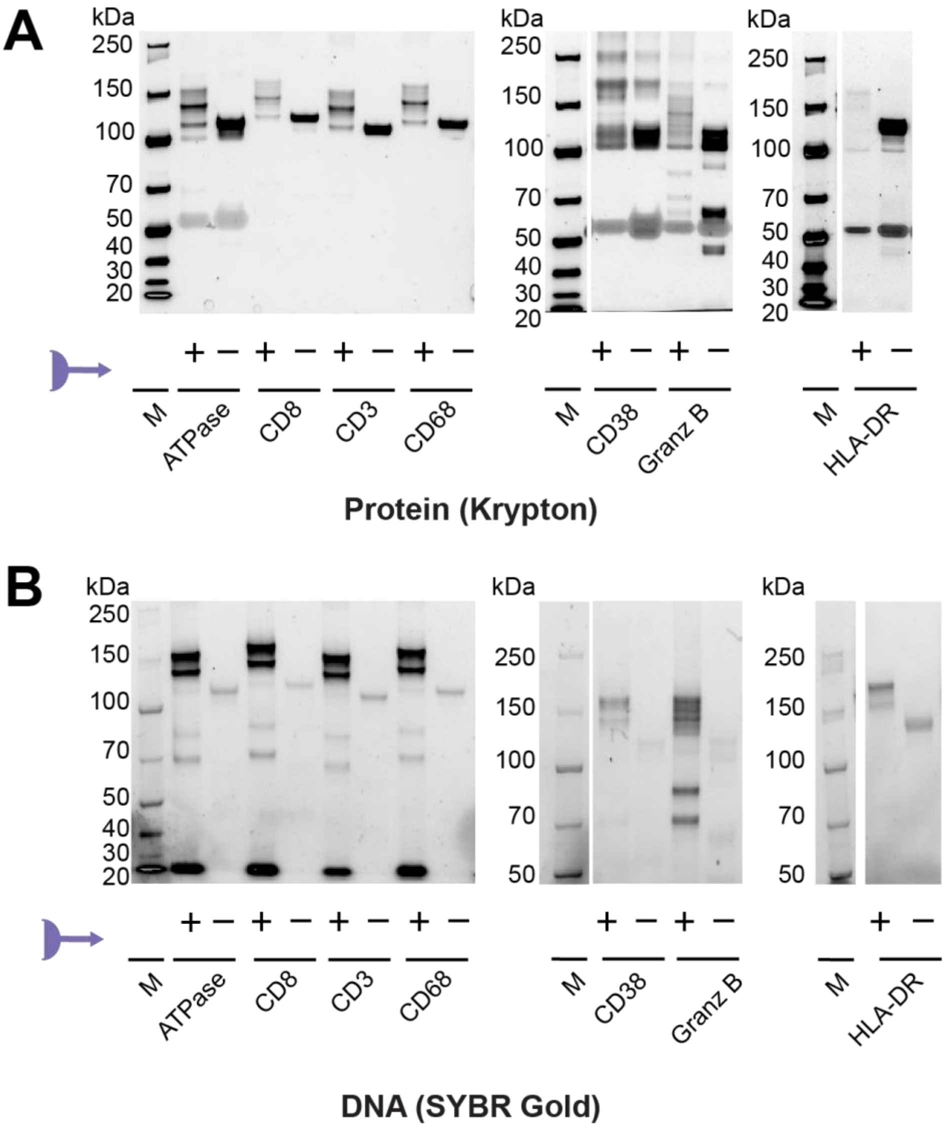


**Supplementary Figure S10. Non-reducing SDS–PAGE analysis of UnO-mediated barcoding of antibodies**. **(A)** 3-8% Tris-acetate gel that show the UnO-barcoded (+) vs. non-conjugated (-) antibodies targeting Na/K ATPase, CD8, CD3, CD68, CD38, Granzyme B, HLA-DR visualized with Krypton protein stain. **(B)** Same gels visualized with SYBR Gold stain. M: Marker lanes.

**Supplementary Table 1. Overview of Immuno-SABER and SABER-Flex detection schemes**

| Aspect | Immuno-SABER<br>(Saka et al., 2019[1]) | SABER-Flex<br>(based on bDNA amplifiers from Xia et al., 2019[2]) |
|---|---|---|
| ssDNA probe (concatemer) synthesis method | Primer Exchange Reaction (Kishi et al., 2018[3]) | Oligonucleotide synthesis (ssDNA) or PCR, *in vitro* transcription (IVT), reverse transcription amplification, or lambda DNAse treatment of dsDNA matrix |
| Probe synthesis yield (per 1 enzymatic reaction) | Low (2-10 µg) | High (100-500 µg, IVT) |
| Hands-on synthesis time | ~0.5–1 h | 0 (no IVT); ~4.5–6 h (w/ IVT) |
| Typical probe length, nt | 300-700 | ≤200 |
| Signal amplification | Yes | Yes (with branching) |
| Branching (secondary probe) | Optional | Recommended |
| Plexity | Up to 50 | 16 |

**Supplementary Table 2. List of antibodies, protein G barcodes, SABER-Flex concatemers and the respective imagers.** "Antibody-barcode" column indicates antibody and the respective Immuno-SABER barcode combination (as previously described[1], here shortened to the last 32 nucleotides instead of the original 42 nucleotide barcodes) conjugated through Protein G. In "Primary concatemer" column, binding sequences of the primary concatemers are complementary to the cognate SABER barcodes (bc1, bc4, bc5) and are indicated in bold. In "Secondary concatemer" column, the binding sequences are complementary to 5x20-mer repeats of the primary concatemers and are indicated in **bold**. In the "Imager" column, the binding sequences are complementary to 5x15-mer repeats of the secondary concatemer. See SABER-Flex scheme in **Fig. 1C** of the main text for reference. Underlined - PCR amplification handles[2]. For antibody information see **Supplementary Table 6.**

| Antibody -barcode | SABER barcode (5′–3′) | Primary concatemer (5′–3′) | Secondary concatemer (5′–3′) | Imager (5′–3′) |
|---|---|---|---|---|
| CD3-bc1 | /AmC6/ttCCGCCAAATCTCCGTGTCCTTAACCGACCTAT | GCCCCATCATGTGCCTTTCC (ACACTTTCACCTTCCCATTAtt)x5 **ATAGGTCGGTTAAGGACACGGAGATTTGGCGG** TGGATTTCCCTCTCGCAGGC | GCCCCATCATGTGCCTTTCC (AACTACCTCTAGGACtt)x5 **TAATGGGAAGGTGAAAGTGT** TGGATTTCCCTCTCGCAGGC | /ATTO 565/tt-GTCCTAGAGGTAGTT |
| CD8-bc4 | /AmC6/ttCTTCGCGTGTTGTCTCGTCTGGGTATTGCGTT | GCCCCATCATGTGCCTTTCC (TCCCAACACATCCTATCTCAtt)x5 **AACGCAATACCCAGACGAGACAACACGCGAAG** TGGATTTCCCTCTCGCAGGC | GCCCCATCATGTGCCTTTCC (AATCGCGACCCTACAtt)x5 **TGAGATAGGATGTGTTGGGA** TGGATTTCCCTCTCGCAGGC | /Alexa Fluor 488/tt-TGTAGGGTCGCGATT |
| CD68-bc5 | /AmC6/ttCGATCCTACCCTTAAAGTTACTGCGCACCCT | GCCCCATCATGTGCCTTTCC (ACCCATTACTCCATTACCATtt)x5 **AGGGTGCGCAGTAACTTTAAGGGTAGGATCG** TGGATTTCCCTCTCGCAGGC | GCCCCATCATGTGCCTTTCC (ATAGGCGTACGAGGGtt)x5 **ATGGTAATGGAGTAATGGGT** TGGATTTCCCTCTCGCAGGC | /Alexa Fluor 647/tt-CCCTCGTACGCCTAT |

**Supplementary Table 3. List of Immuno-SABER barcode sequences.**

All barcode oligos were purchased from IDT at 100 nmol synthesis scale with HPLC purification.

| Barcode | Name | Oligo sequence (5′–3′) |
|---|---|---|
| 0 | saber-c*-u*-b0 | CTTGCTGTTCTTCC-tt-CTGAGACTGGATGC-tt-AATTCTATGACACCGCCACGCCCTATATCC |
| 1 | saber-c*-u*-b1 | CTTGCTGTTCTTCC-tt-CTGAGACTGGATGC-tt-ATTATCCCTACCGCCAAATCTCCGTGTCCT |
| 2 | saber-u*-b2 | CTGAGACTGGATGC-tt-CGTTATCGCCGCCTTATCCACTGTACGATC |
| 3 | saber-c*-u*-b3 | CTTGCTGTTCTTCC-tt-CTGAGACTGGATGC-tt-GTTTCCTATATTTAGCGTCCGTGTCGTTCT |
| 4 | saber-c*-u*-b4 | CTTGCTGTTCTTCC-tt-CTGAGACTGGATGC-tt-TATCTTAAGTCTTCGCGTGTTGTCTCGTCT |
| 8 | saber-c*-u*-b8 | CTTGCTGTTCTTCC-tt-CTGAGACTGGATGC-tt-AACAATTCAGCTCCGCCTTATACCGTCTTA |
| 10 | saber-c*-u*-b10 | CTTGCTGTTCTTCC-tt-CTGAGACTGGATGC-tt-GTCCTCGCTCTTTCCGCATTTTCCCGTATG |
| 11 | saber-c*-u*-b11 | CTGAGACTGGATGC tt TGTCTAAATTCTAATGCCGCCCTATGCCGC |
| 12 | saber-u*-b12 | CTGAGACTGGATGC-tt-CCTTCCGCCGTATGAATTTGACCCGAAGCC |
| 14 | saber-c*-u*-b14 | CTTGCTGTTCTTCC-tt-CTGAGACTGGATGC-tt-CCAACCTCTCGTACCAAATTCCGCCACTCA |

***Note:*** u* (CTGAGACTGGATGC) denotes a common barcoding sequence, complementary to cnvK-modified universal oligo (GCATCCA[cnvk]TCTCAG) in UnO. Underlined T denotes cnvK acceptor nucleotide base. b0, b1, … denote barcode sequences following the previous numbering scheme for Immuno-SABER[1]. c* (CTTGCTGTTCTTCC) denotes an optional common capture sequence that potentially can be used for pull-down of barcoded antibodies. This optional domain was not included in barcode 12.

**Supplementary Table 4. List of Immuno-SABER primers and corresponding PER conditions.**

| Barcode | Name | Oligo sequence (5′–3′) | Hairpin ID | Hairpin sequence | Hairpin concent-ration | Time | Imager |
|---|---|---|---|---|---|---|---|
| 0 | saber-b0*-p38.38 | GGA TAT AGG GCG TGG CGG TGT CAT AGA ATT ttt AACATACTAaAACATACTA | h.38.38 | AAACATACTAGGGCCTTTTGGCCCTAGTATGTTTTAGTATGTT/3InvdT/ | 5 µM | 2 h | /ATTO 565/tt-TAGTATGTT-t-TAGTATGTT-t/3InvdT/ |
| 1 | saber-b1*.p28.28 | AGG ACA CGG AGA TTT GGC GGT AGG GAT AAT ttt CAACTTAACaCAACTTAAC | h.28.28 | ACAACTTAACGGGCCTTTTGGCCCGTTAAGTTGTGTTAAGTTG/3InvdT/ | 3 µM | 2 h | /ATTO 488/tt-GTTAAGTTG-t-GTTAAGTTG-t/3InvdT/ |
| 2 | saber-b2*.p30.30 | GATCGTACAGTGGATAAGGCGGCGATAACG ttt AATACTCTCaAATACTCTC | h.30.30 | AAATACTCTCGGGCCTTTTGGCCCGAGAGTATTTGAGAGTATT/3InvdT/ | 5 µM | 2 h | /ATTO 565/tt-GAGAGTATT-t-GAGAGTATT-t/3InvdT/ |
| 3 | saber-b3*.p27.27 | AGA ACG ACA CGG ACG CTA AAT ATA GGA AAC ttt CATCATCATaCATCATCAT | h.27.27 | ACATCATCATGGGCCTTTTGGCCCATGATGATGTATGATGATG/3InvdT/ | 0.75 µM | 3 h | /ATTO 565/tt-ATGATGATG-t-ATGATGATG-t 3InvdT/ |
| 4 | saber-b4*-p37.37 | AGA CGA GAC AAC ACG CGA AGA CTT AAG ATA ttt TTTCTCTTCaTTTCTCTTC | h.37.37 | ATTTCTCTTCGGGCCTTTTGGCCCGAAGAGAAATGAAGAGAAA/3InvdT/ | 8.5 µM | 2 h | /Alexa Fluor 647/tt-GAAGAGAAA-t-GAAGAGAAA-t/3InvdT/ |
| 8 | saber-b8*-p25.25 | TAA GAC GGT ATA AGG CGG AGC TGA ATT GTT ttt CCAATAATAaCCAATAATA | h.25.25 | ACCAATAATAGGGCCTTTTGGCCCTATTATTGGTTATTATTGG/3InvdT/ | 1.5 µM | 3 h | /ATTO 488/tt-TATTATTGG-t -ATTATTGG-t/3InvdT/ |
| 10 | saber-b10*.p42.42 | CAT ACG GGA AAA TGC GGA AAG AGC GAG GAC ttt CTTACAAACaCTTACAAAC | h.42.42 | ACTTACAAACGGGCCTTTTGGCCCGTTTGTAAGTGTTTGTAAG/3InvdT/ | 5 µM | 2 h | /ATTO 488/tt-GTTTGTAAG-t-GTTTGTAAG-t/3InvdT/ |
| 11 | saber-b11*.p26.26 | GCGGCATAGGGCGGCATTAGAATTTAGACA ttt ATAAACCTAaATAAACCTA | h.26.26 | AATAAACCTAGGGCCTTTTGGCCCTAGGTTTATTTAGGTTTAT/3InvdT/ | 9 µM | 3 h | /ATTO 550/tt-TAGGTTTAT-t-TAGGTTTAT-t/3InvdT/ |
| 12 | saber-b12*.p | GGCTTCGGGTCAAATTCATACG | h.43.43 | AACAAATAACGGGCCTT | 5 µM | 2 h | /Alexa Fluor 647/tt- |

| | | | | | | | |
|---|---|---|---|---|---|---|---|
| | 43.43 | GCGGAAGG ttt ACAAATAACaAC AAATAAC | | TTGGCCCG TTATTTGTT GTTATTTGT /3InvdT/ | | | GTTATTTGT-t- GTTATTTGT-t /3InvdT/ |
| 14 | saber-b14*.p 41.41 | TGA GTG GCG GAA TTT GGT ACG AGA GGT TGG ttt CAATCAAAAaCA ATCAAAA | h.41.41 | ACAATCAAA AGGGCCTT TTGGCCCT TTTGAT TGTTTTTGA TTG/3InvdT/ | 4.5 µM | 3 h | /Alexa Fluor 647/ tt-TTTTGATTG-t-TTTTGATTG-t /3InvdT/ |

***Note***: b0*, b1*, … denote binding sequences of concatemers, complementary to the cognate antibody barcodes (b0, b1, …). p25.25, p26.26, … denote direct 20-mer repeats of the respective PER primer sequences. See *Saka et al., 2019* for reference.

**Supplementary Table 5. List of Immuno-SABER antibodies, respective barcodes and primers.**

| Bar-code | Cycle # | Antibody | Dilution used | Vendor | Clone | Catalog # | Isotype | Primer | Fluor |
|---|---|---|---|---|---|---|---|---|---|
| saber-c*-u*-b0 | 1 | alpha-Tubulin | 1:500 | Abcam | EP1332Y | ab52866 | rbIgG | saber-b0*-p38.38 | ATTO 565 |
| saber-c*-u*-b1 | 1 | SP100 | 1:200 | Atlas Antibodies | Polyclonal | HPA016707 | rbIgG | saber-b1*-p28.28 | ATTO 488 |
| saber-u*-b2 | 2 | SON | 1:100 | Atlas Antibodies | polyclonal | HPA031755 | rbIgG | saber-b2*.p30x2 | ATTO 565 |
| saber-c*-u*-b3 | 1 | G3BP1 | 1:100 | Cell Signaling Technology | E9G1M | 61559T | rbIgG | saber-b3*-p27.27 | ATTO 565 |
| saber-c*-u*-b4 | 2 | Coilin | 1:100 | Cell Signaling Technology | D2L3J | 14168T | rbIgG | saber-b4*--p37.37 | Alexa Fluor 647 |
| saber-c*-u*-b8 | 3 | Vimentin | 1:100 | Cell Signaling Technology | D21H3 | 5741T | rbIgG | saber-b8*-p25.25 | ATTO 488 |
| saber-c*-u*-b10 | 2 | TFAM | 1:100 | Thermo | 18G102B2E11 | MA5-16148 | rbIgG2b | saber-b10*-p42.42 | ATTO 488 |
| saber-c*-u*-b11 | 3 | TOMM20 | 1:100 | Sigma | 4F3 | WH0009804M1-100UG | mIgG1 | saber-b11*-p26x2 | ATTO 565 |
| saber-u*-b12 | 4 | SC35 (SRSF2) | 1:100 | Abcam | SC-35 | ab11826 | mIgG1 | saber-b12*-p43x2 | Alexa Fluor 647 |
| saber-c*-u*-b14 | 3 | GM130 | 1:100 | Cell Signaling Technology | D6B1 | 12480T | rbIgG | saber-b14*-p41.41 | Alexa Fluor 647 |

***Note:*** in “Isotype” column “m” or “rb” before IgG stand for mouse or rabbit origin, respectively.

**Supplementary Table 6. List of CODEX antibodies and respective barcodes.**

| Barcode | Sequence (5′–3′) | Antibody target | Clone | Manufacturer | Catalog # | Fluorophore |
|---|---|---|---|---|---|---|
| u*_cod_2 | CTGAGACTGGATGCtttATGGTTTAGGACTAC | CD38 | EPR4106 | Abcam | ab108403 | ATTO550 |
| u*_cod_8 | CTGAGACTGGATGCtttCGCAGATGAATATTC | CD8 | D8A8Y | Cell Signaling Technology | 81575SF | ATTO647 |
| u*_cod_41 | CTGAGACTGGATGCtttTGTATGAGTAGTAATCT | CD68 | D4B9C | Cell Signaling Technology | 26042SF | ATTO550 |
| u*_cod_65 | CTGAGACTGGATGCtttGATAAATATTTTACAGAGT | HLA DR | EPR3692 | Abcam | ab215985 | ATTO550 |
| u*_cod_67 | CTGAGACTGGATGCtttGACGACGAAGGC | Na/K ATPase | EP1845Y | Abcam | ab283340 | ATTO647 |
| u*_cod_77 | CTGAGACTGGATGCtttATTTCAACAAATATTGTT | CD3 | D7A6E | Cell Signaling Technology | 24581SF | ATTO647 |
| u*_cod_81 | CTGAGACTGGATGCtttCAAGGAACTACCGA | Granzyme B | EPR20129-217 | Abcam | ab219803 | ATTO647 |

### Supplementary References


1. Saka, S.K., Wang, Y., Kishi, J.Y., Zhu, A., Zeng, Y., Xie, W., Kirli, K., Yapp, C., Cicconet, M., Beliveau, B.J., et al. (2019). Immuno-SABER enables highly multiplexed and amplified protein imaging in tissues. Nat. Biotechnol. *37*, 1080–1090. https://doi.org/10.1038/s41587-019-0207-y.

2. Xia, C., Babcock, H.P., Moffitt, J.R., and Zhuang, X. (2019). Multiplexed detection of RNA using MERFISH and branched DNA amplification. Sci. Rep. *9*, 7721. https://doi.org/10.1038/s41598-019-43943-8.

3. Kishi, J.Y., Schaus, T.E., Gopalkrishnan, N., Xuan, F., and Yin, P. (2018). Programmable autonomous synthesis of single-stranded DNA. Nat. Chem. *10*, 155–164. https://doi.org/10.1038/nchem.2872.
